# Spatially compartmentalized phase regulation of a Ca^2+^-cAMP-PKA oscillatory circuit

**DOI:** 10.1101/2020.01.10.902312

**Authors:** Brian Tenner, Michael Getz, Brian Ross, Donya Ohadi, Christopher H. Bohrer, Eric Greenwald, Sohum Mehta, Jie Xiao, Padmini Rangamani, Jin Zhang

**Affiliations:** Department of Biophysics and Biophysical Chemistry, The Johns Hopkins University School of Medicine, Baltimore, MD 21505, USA.; Department of Pharmacology, University of California, San Diego, La Jolla, CA 92093, USA.; Chemical Engineering Graduate Program, University of California, San Diego, La Jolla, CA 92093, USA.; Department of Mechanical and Aerospace Engineering, University of California, San Diego, La Jolla, CA 92093, USA.

## Abstract

Signaling networks are spatiotemporally organized in order to sense diverse inputs, process information, and carry out specific cellular tasks. In pancreatic β cells, Ca^2+^, cyclic adenosine monophosphate (cAMP), and Protein Kinase A (PKA) exist in an oscillatory circuit characterized by a high degree of feedback, which allows for specific signaling controls based on the oscillation frequencies. Here, we describe a novel mode of regulation within this circuit involving a spatial dependence of the relative phase between cAMP, PKA, and Ca^2+^. We show that nanodomain clustering of Ca^2+^-sensitive adenylyl cyclases drives oscillations of local cAMP levels to be precisely in-phase with Ca^2+^ oscillations, whereas Ca^2+^-sensitive phosphodiesterases maintain out-of-phase oscillations outside of the nanodomain, representing a striking example and novel mechanism of cAMP compartmentation. Disruption of this precise in-phase relationship perturbs Ca^2+^ oscillations, suggesting that the relative phase within an oscillatory circuit can encode specific functional information. This example of a signaling nanodomain utilized for localized tuning of an oscillatory circuit has broad implications for the spatiotemporal regulation of signaling networks.

## INTRODUCTION

Cyclic adenosine monophosphate (cAMP) and Ca^2+^ act as essential second messengers in almost every cell type and regulate many functional pathways within a cell, such as hormonal signal transduction, metabolism, and secretion (Sassone-Corsi 2012; Clapham 2007). In some cell types, including neurons, cardiomyocytes, and pancreatic β cells, these messengers’ concentrations oscillate intracellularly (Dupont et al. 2011; Dyachok et al. 2006.), and the oscillations encode critical signaling information (e.g. signal strength, duration, and target diversity) into parameters such as frequency and amplitude (Berridge et al. 1998; De Pitta et al. 2008; Parekh 2010). This is perhaps best exemplified in the β cell where oscillations of Ca^2+^ drive pulsatile insulin secretion (Rorsman et al. 2018) as well as oscillations in cAMP levels (Tengholm 2012; Nesher et al. 2002). Furthermore, Ca^2+^, cAMP, and the downstream cAMP-dependent kinase Protein Kinase A (PKA) constitute a highly-coordinated oscillatory circuit responsible for integrating metabolic and signaling information (Ni et al. 2011). In addition to temporal control, biochemical pathways are also spatially organized within the cell (White et al. 2005; Smith et al. 2002). Both Ca^2+^ and cAMP are highly spatially compartmentalized and form signaling microdomains or nanodomains (Peterson 2002; Calebiro et al. 2014). While Ca^2+^ levels are locally controlled by channels, pumps, and intracellular buffering systems (Clapham 2007; Stern 1992), cAMP is thought to be regulated via controlled synthesis by adenylyl cyclases (ACs) and degradation by phosphodiesterases (PDEs) (Hanoune et al. 2001; Bender et al. 2006). Despite extensive studies on cAMP compartmentation, the mechanisms that spatially constrain this mobile second messenger remain poorly understood (Lohse et al. 2017; Musheshe et al. 2018; Saucerman et al. 2013). Furthermore, it is not clear how spatial regulation of a second messenger influences its dynamic behaviors in the context of coordinated oscillations.

In this study, we investigated the spatiotemporal organization of the Ca^2+^-cAMP-PKA oscillatory circuit in pancreatic β cells and discovered that the relative, oscillatory phase between cAMP/PKA and Ca^2+^ is spatially regulated within signaling nanodomains. By combining live-cell dynamic imaging, super-resolution microscopy, and computational modeling, we further found that fine-scale, compartment-specific perturbations of this precise phase dynamic impacts Ca^2+^ oscillations in the β cells. These findings suggest that the relative phase in oscillatory signaling circuits, like the amplitude and frequency, represents yet another mode of informational encoding and processing, which is subjected to spatiotemporal regulation within the cell.

## RESULTS

### The relative phase of β cell cAMP and Ca^2+^ oscillations is compartmentalized

In order to study the spatiotemporal relationship between key players of the Ca^2+^-cAMP-PKA circuit, we chose to focus our attention on an important class of molecular scaffolds, A-Kinase Anchoring Proteins (AKAPs), which are responsible for recruiting PKA to specific substrates at distinct subcellular locations. In several excitable cell types, the plasma membrane (PM) localized scaffold protein AKAP79 (AKAP150 rodent ortholog) has been shown to organize a macromolecular complex with binding partners that include PKA, the voltage-gated Ca^2+^ channel Ca_V_1.2, Protein Kinase C (PKC), the Ca^2+^/calmodulin-dependent protein phosphatase calcineurin, Ca^2+^-sensitive ACs, AMPA receptors, and many others (Gold et al. 2011). Due to the extensive and multivalent nature of AKAP79/150 and a report describing the functional impairment of glucose-stimulated insulin secretion (GSIS) in pancreatic β cells upon its knock-out (Hinke et al. 2012), we hypothesized that the AKAP79/150 scaffold might play an important role in the spatiotemporal regulation of the Ca^2+^-cAMP-PKA oscillatory circuit. Specifically, we were interested in testing if AKAP79/150 is able to create a spatially-distinct compartment in which recruitment of signaling effectors can locally fine-tune and reshape signaling dynamics within the circuit (Beene et al. 2007; Greenwald et al. 2011). In order to test this hypothesis, we monitored intracellular cAMP and Ca^2+^ using the FRET-based cAMP biosensor (Ci/Ce)Epac2-camps (Everett et al. 2013) and the red Ca^2+^ indicator RCaMP (Akerboom et al. 2013) in MIN6 β cells. By fusing (Ci/Ce)Epac2-camps to the full-length AKAP79 scaffold and transiently transfecting the targeted sensor, we measured cAMP concentration changes in the immediate vicinity of AKAP79/150 (**Fig. 1a**). As a control, we also targeted the cAMP probe to the general plasma membrane by adding a lipid modification domain (Wachten et al. 2010). These targeted biosensors allowed us to compare the dynamics within the AKAP79/150-specific compartment versus the general plasma membrane compartment (**Fig. 1a**).

**Figure 1.**
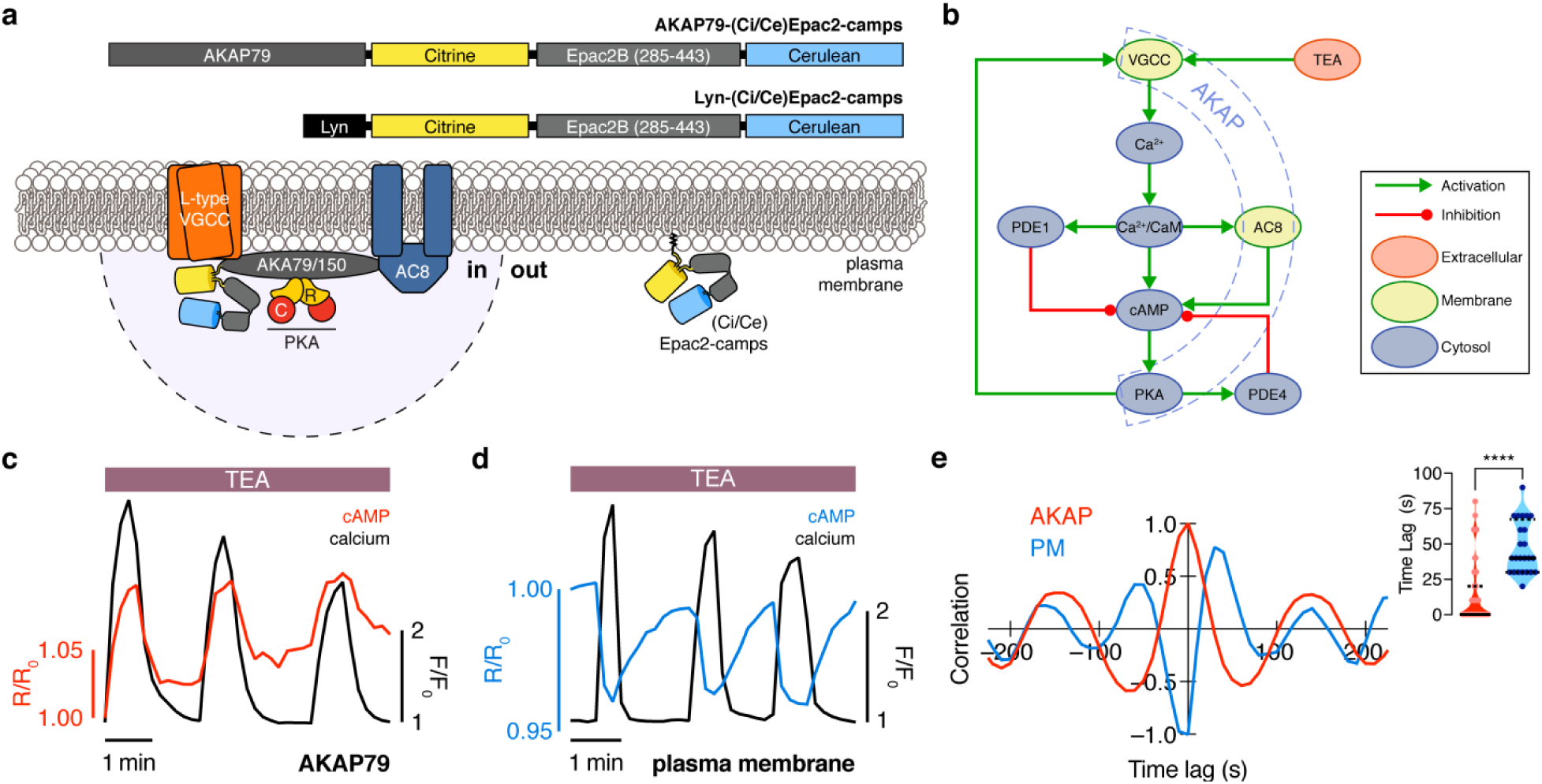
The phase of oscillating cAMP is shifted between the AKAP79/150 compartment and the general plasma membrane compartment, relative to Ca^2+^. (a) Depiction of the AKAP79 compartment and plasma membrane compartment, including the targeted cAMP biosensor (Ci/Ce)Epac2-camps to measure the compartment-specific cAMP signaling. Schematics of the lyn-(Ci/Ce)Epac2-camps and AKAP79-(Ci/Ce)Epac2-camps sensors. (b) Network diagram describing the key players in the Ca^2+^-cAMP-PKA oscillatory circuit in the β cell. (c) Representative single cell trace of an in-phase oscillating β cell with AKAP79-(Ci/Ce)Epac2-camps and RCaMP, whole-cell fluorescence measured. Red trace is cAMP (cyan direct channel divided by CY-FRET channel) and black trace is Ca^2+^ (RFP). (d) Representative single cell trace of an out-of-phase oscillating β cell with lyn-(Ci/Ce)Epac2-camps and RCaMP, whole-cell fluorescence measured. Blue trace is cAMP (cyan direct channel divided by CY-FRET channel) and black trace is Ca^2+^ (RFP). (e) Cross-correlation between the oscillatory Ca^2+^ and cAMP signals from the representative in-phase AKAP79 (red) and out-of-phase plasma membrane (PM, blue) β cells from c, d. Time lag (sec) between the cAMP and Ca^2+^ signals for the two compartments (AKAP79/150, red, is 13 ± 3 sec n = 60 and PM, blue, is 47 ± 4 sec n = 24) (p<0.05).

Although both targeted sensors were trafficked to and distributed along the plasma membrane (**Extended Fig. 1**), we observed notable differences in their respective cAMP signals relative to Ca^2+^ oscillations after triggering the circuit (**Fig. 1b**) with tetraethylammonium chloride (TEA, 20mM), a potent K^+^ channel blocker. cAMP oscillations measured within the AKAP79/150 compartment were in-phase with oscillating Ca^2+^ such that each transient spike in intracellular Ca^2+^ was closely associated with a transient increase in cAMP (**Fig. 1c**) (n = 60 cells). This was in sharp contrast to cAMP oscillations measured within the general plasma membrane compartment where each local Ca^2+^ peak corresponded to a local trough in cAMP (n = 24), followed by a slow reversal of both signals to a pre-stimulated baseline (**Fig. 1d**). While these out-of-phase cAMP-Ca^2+^ oscillations were consistent with those observed in the cytoplasm of β cells, in phase cAMP-Ca^2+^ oscillations had not be observed under these conditions (Ni et al. 2011; Landa et al. 2005). To quantify the cAMP-Ca^2+^ phase relationship, we measured the lag time by calculating the cross-correlation between the two normalized, oscillatory signals and finding the shortest delay yielding the maximum correlation (see Supplementary Information for details) (**Fig. 1e**). In-phase cAMP oscillations corresponded to short lag times (typically <20 sec) while out-of-phase oscillations mostly possessed longer lag times. Within the AKAP79/150 compartment, cAMP lagged behind Ca^2+^ by an average of only 13 ± 3 sec (n = 60); however, cAMP within the general plasma membrane compartment oscillated with a lag time of 47 ± 4 sec (n = 24), relative to Ca^2+^ (**Fig. 1e**). This stark difference in the cAMP-Ca^2+^ phase relationship suggests that the relative phase of this oscillatory circuit is compartmentalized and hints at differential regulation of the circuit between the AKAP79/150 compartment and the general plasma membrane compartment.

### Oscillatory phase is regulated by balanced activities of Ca^2+^-sensitive ACs and PDEs

Given that in-phase cAMP oscillations were only observed within the AKAP79/150 compartment (**Fig. 1c**) and out-of-phase cAMP oscillations were observed in the general plasma membrane compartment (**Fig. 1d**) and cytoplasm (Ni et al. 2011), we hypothesized that Ca^2+^ oscillations are coupled to cAMP oscillations by a ubiquitous mechanism throughout the cell, while additional mechanisms specifically regulate the phase relationship within the AKAP79/150 compartment. We first sought to identify the component that is responsible for coupling cAMP dynamics to Ca^2+^ dynamics globally. Since TEA induces continuous Ca^2+^ oscillations, we determined the temporal relationship between Ca^2+^ and cAMP at the general plasma membrane more precisely by measuring the impulse response of the circuit following a transient membrane depolarization. After the addition of KCl (15 mM) followed by a subsequent washout to elicit a transient influx of Ca^2+^ (Dou et al. 2014), we observed a synchronous cAMP decrease (n = 20) followed by a return to baseline (**Fig. 2a**). This data suggests that increasing cytosolic Ca^2+^ was coupled to a decrease in cAMP at the plasma membrane via Ca^2+^-sensitive AC or PDE activities. Given that Ca^2+^-inhibited ACs (AC5, AC6) have low specific activity both in the presence and absence of physiological Ca^2+^, as well as relatively low expression in the pancreas (Defer et al. 2000), we instead focused on probing the roles of PDEs. The Ca^2+^-dependent PDE1 family in MIN6 cells, specifically PDE1C, has been implicated in modulating GSIS (Han et al. 1999). Indeed, acute addition of 8MM-IBMX (100 μM), a relatively selective PDE1 inhibitor, effectively uncoupled cAMP dynamics from Ca^2+^ oscillations (**Fig. 2b**, **Extended Fig. 2a**) (n = 18), indicating that Ca^2+^-triggered activation of PDE1 mediates the transient cAMP decreases. We also observed that the overall increase in cAMP led to an increase in the Ca^2+^ oscillation frequency, consistent with the previously identified role of cAMP/PKA in regulating the Ca^2+^ oscillations (Ni et al. 2011). We tested the roles of two additional families of abundant PDEs in pancreatic β cells, PDE3 and PDE4, by acute pharmacologic inhibition. While treating cells with either milrinone (PDE3 inhibitor, 10 μM, n = 12) or rolipram (PDE4 inhibitor, 1 μM, n = 15) slightly increased cAMP levels, neither inhibitor had an effect on cAMP-Ca^2+^ coupling or relative phase (**Extended Fig. 2b,c**). These data suggest that PDE1 is the key component that couples Ca^2+^ and cAMP oscillations within this signaling circuit.

**Figure 2.**
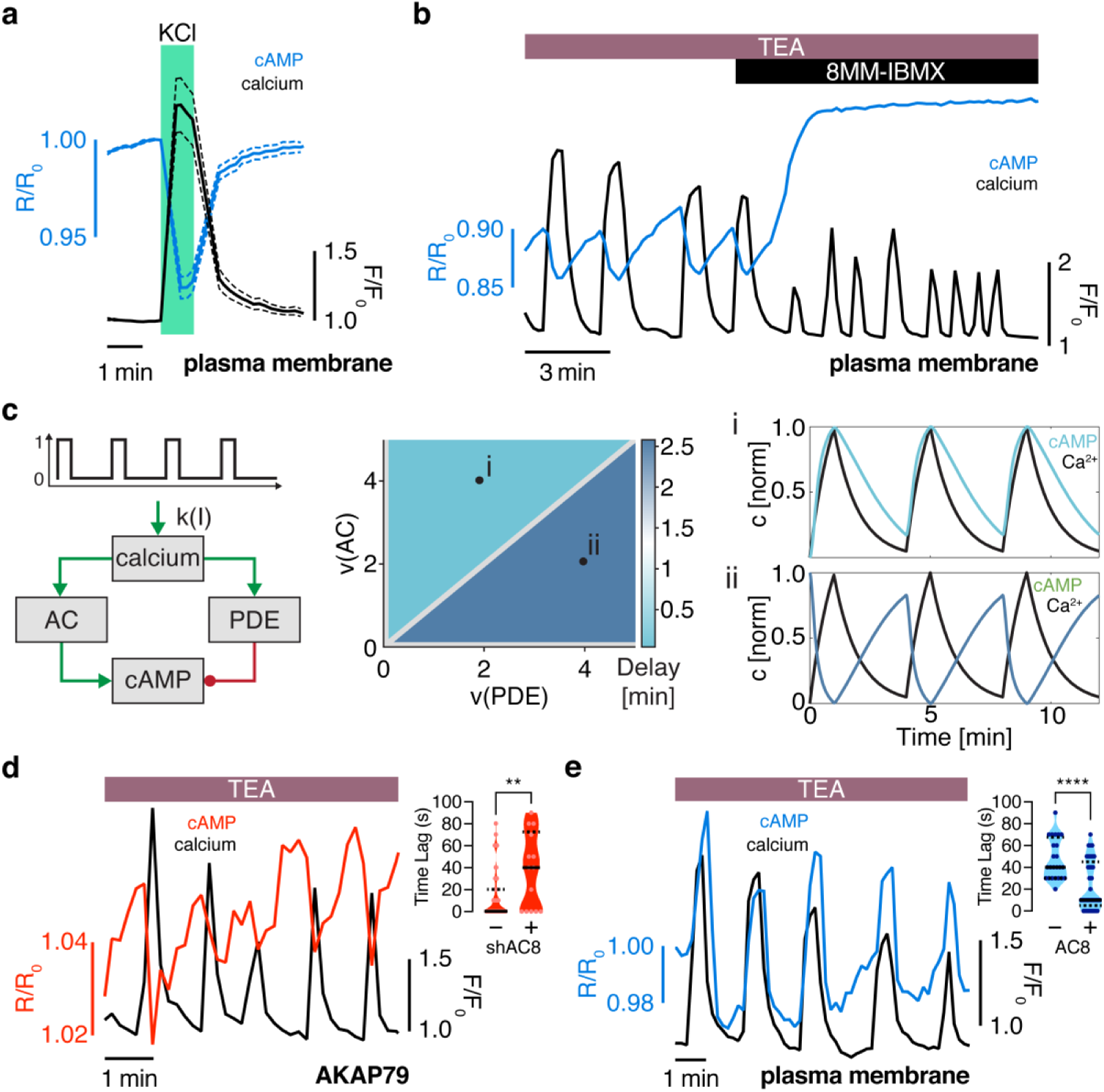
The oscillation phase is regulated by a balance between Ca^2+^-sensitive AC and PDE activity. (a) Impulse response of plasma membrane cAMP (blue) to a spike in Ca^2+^ entry (black), triggered by KCl-mediated membrane depolarization (wash in/out). The transient decrease in PM-cAMP is coupled to the transient increase in intracellular Ca^2+^. (b) Acute inhibition of Ca^2+^-sensitive PDE1 decouples the out-of-phase PM-cAMP oscillations from Ca^2+^ oscillations, as observed in this representative cell trace (Ca^2+^ – black, PM-cAMP – blue). (c) The oscillatory phase of cAMP can be manipulated by tuning the relative activity of Ca^2+^-sensitive PDE and AC, as demonstrated by a simplified model. (d) Knocking down AC8 is correlated with an increase in the time lag for oscillatory cAMP at the AKAP79/150 microdomain (37 ± 9 sec, n = 11), indicating more cells exhibiting out-of-phase cAMP oscillations (representative cell trace, Ca^2+^ – black, AKAP79/150-cAMP – red). (e) Co-expressing AC8 is sufficient to reverse the phase at the PM to in-phase (23 ± 2 sec, n = 56) (representative cell trace, Ca^2+^ – black, PM cAMP – blue).

How is the phase relationship between Ca^2+^ and cAMP regulated within distinct signaling compartments? In order to gain a more quantitative understanding of the regulation of the cAMP-Ca^2+^ phase relationship, we created a simplified mathematical model involving Ca^2+^, cAMP, and Ca^2+^-driven PDE and AC activity components (Cooper et al. 1995) (**Fig. 2c**, see Supplementary Material for details). This simple circuit represents the key aspects of the oscillatory cAMP-Ca^2+^ circuit and is applicable to different signaling compartments. Opposite to the Ca^2+^-stimulated PDE1 (Ang et al. 2002) is the Ca^2+^-stimulated AC8 (Masada et al. 2008; Masada et al. 2012), an abundant Ca^2+^-sensitive transmembrane AC isoform in β cells that has been shown to mediate sustained insulin secretion and associate with the AKAP79/150 scaffold (Dou et al. 2014; Willoughby et al. 2010; Willoughby et al. 2006). By computationally manipulating the activity of each arm, we found that cAMP can oscillate either out-of-phase or in-phase when a simulated Ca^2+^ pulse train is used as an input (**Fig. 2c**). In particular, when the relative activity of PDE1 is greater than the activity of AC8, Ca^2+^-driven cAMP degradation dominates, resulting in an out-of-phase cAMP-Ca^2+^ relationship. On the other hand, if the relative activity of AC8 is greater than that of PDE1, Ca^2+^-stimulated cAMP production is favored and an in-phase relationship is observed, consistent with previous modeling studies (Peercy et al. 2015; Fridlyand et al. 2007).

Thus, our simplified model indicates that the phase relationship can be tuned by altering the relative strength between Ca^2+^-sensitive ACs and PDEs (**Fig. 2c**). This model provided a blueprint for understanding the interplay between the Ca^2+^-stimulated AC/PDE balance and the cAMP-Ca^2+^ phase relationship within the AKAP79/150 compartment. Based on the findings from our model, we predicted that decreasing the relative contribution of AC8 will shift the cAMP-Ca^2+^ phase relationship from in-phase to out-of-phase. To test this prediction, we knocked down endogenous AC8 in the MIN6 cells as previously done (Raoux et al. 2015) and observed that most cells exhibited an out-of-phase cAMP oscillation within the AKAP79/150 compartment (average lag time 37 ± 9sec, n = 11) (**Fig. 2d**), indicating an AC8-specific role in mediating the cAMP-Ca^2+^ phase signature.

Conversely, increasing the relative contribution of AC8, for example by increasing the concentration of AC8 throughout the plasma membrane, should shift the cAMP-Ca^2+^ phase relationship from out-of-phase to in-phase. To test this prediction, we overexpressed full-length AC8 and examined the effect in the general plasma membrane compartment. Interestingly, we found that AC8 overexpression reversed the out-of-phase cAMP-Ca^2+^ phase relationship in a titratable manner where the percentage of in-phase oscillating cells correlated with increasing amounts of the co-transfected AC8 (average lag time 23 ± 2 sec, n = 56) (**Fig. 2e**, **Extended Fig. 3a-c**). This data demonstrates that higher levels of AC8 are sufficient to reverse the cAMP phase at the plasma membrane. In summary, these phase manipulation experiments suggest that the cAMP-Ca^2+^ phase relationship is representative of a sensitive, compartmentalized balance between the Ca^2+^-stimulated activities of PDE1 and AC8.

### Membrane-localized AKAP150:AC8 nanoclusters regulate cAMP-Ca^2+^ oscillatory phase

The close spatial juxtaposition between the AKAP79/150 and general plasma membrane compartments presents a significant challenge for cAMP compartmentation, in that cAMP oscillations must be distinctly regulated within these adjacent signaling domains. Indeed, how cAMP, a rapidly diffusing small molecule, is spatially compartmentalized in cells is not yet clearly understood, especially given the low catalytic efficiency of a single cAMP-producing AC and degrading PDE (Conti et al. 2014; Lohse et al. 2017). Given that AKAP79/150 exists in nanoclusters at the plasma membrane in multiple cell types (Mo et al. 2017; Zhang et al. 2016) and associates with AC8 in β cells (Willoughby et al. 2010), we hypothesized that AC8 could form nanoclusters on the plasma membrane of MIN6 cells and compartmentalize cAMP dynamics. To test this hypothesis, we examined the spatial organization of AC8 and AKAP150 at the membrane using Stochastic Optical Reconstruction Microscopy (STORM). We found that the AKAP150 molecules were organized in clusters with a mean radius of 127 ± 9 nm and an average nearest-neighbor spacing of 313 ± 20nm between cluster centers (n = 20) (**Fig. 3a**), consistent with several recent reports demonstrating AKAP79/150’s tendency to form nanoclusters in other cell types (Zhang et al. 2016; Tajada et al. 2017; Mo et al. 2017; Purkey et al. 2018). Thus, the AKAP79/150 compartment-specific cAMP phase is likely representative of the balanced cAMP generation and degradation within these AKAP clusters.

**Figure 3.**
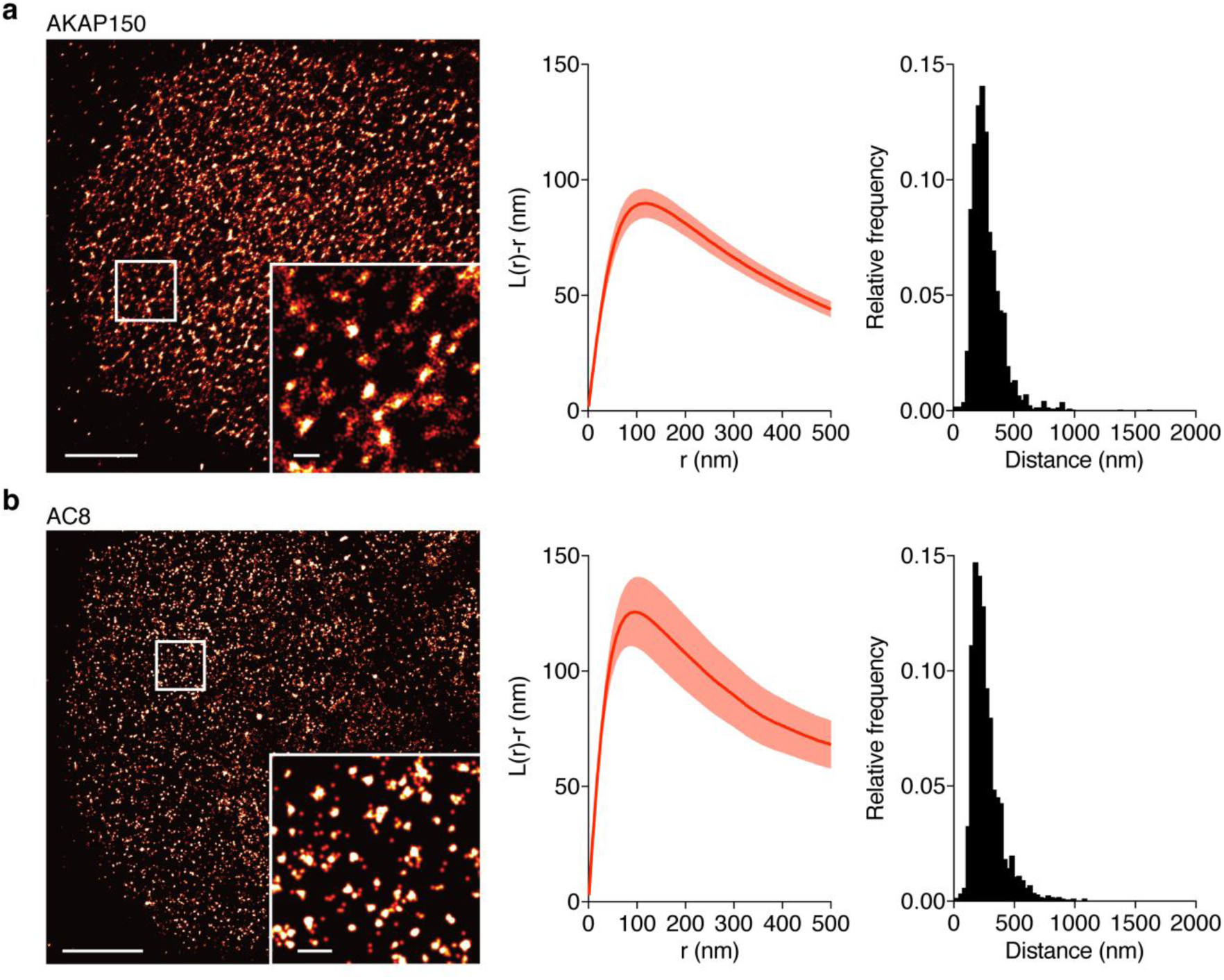
AKAP150 and AC8 both form nanoclusters at the surface of MIN6 β cells. (a) Representative super-resolution STORM image of the AKAP150 scaffold (scale 5μm, inset 500nm). Ripley-K analysis measures the average radii of the nanoclusters and indicates that AKAP150 forms clusters of 127 ± 9 nm, n = 20. The nearest-neighbor distance distribution describes the distance between nanoclusters (average distance for AKAP150 is 313 ± 20 nm). (b) Representative super-resolution STORM image of Ca^2+^-sensitive AC8 (scale 5μm, inset 500nm) depicts AC8 nanoclusters of average radius 88 ± 8 nm and average nearest-neighbor distance 292 ± 16 nm, n = 16.

Due to the known interaction between AKAP79/150 and AC8 (Willoughby et al. 2010), next we probed the spatial organization of AC8. We found AC8 also distributes non-uniformly at the plasma membrane and clusters with a mean radius of 88 ± 8nm and an average nearest-neighbor spacing of 292 ± 16nm between cluster centers (n = 16) (**Fig. 3b**). With the evidence of the nanoscale organization of AKAP150 and AC8 on the plasma membrane, we further hypothesized that the increased spatial density of Ca^2+^-driven cAMP sources within the AKAP150 clusters, in conjunction with dispersed PDE1 in the cytosol (Bender et al. 2006; Goraya et al. 2008), is important in compartmentalizing cAMP production and mediating the in-phase cAMP signal. To test this idea, we sought to build a mathematical framework to describe the spatial compartmentalization of the in- and out-of-phase cAMP-Ca^2+^ oscillations. Briefly, we used the AKAP79/150:AC8 cluster pattern measurements from the STORM imaging to set model parameters in a hexagonal prism domain (200nm edge, 600nm depth) with one AKAP79/150:AC8 cluster centered in the domain for simulation (**Fig. 4a**, see Supplementary Information for model development details). We extended a previous well-mixed β cell model (Ni et al. 2011) to include the Ca^2+^-sensitive PDE1 and a 3D spatial component with cAMP diffusion (*D_cAMP_* = 60 μm^2^/s, Agarwal et al. 2016). By localizing AC8 within the AKAP79/150:AC8 cluster on the plasma membrane face and leaving PDE1 well-mixed throughout the volume, we could simulate Ca^2+^-driven cAMP oscillations that were in-phase within the immediate vicinity of a cluster, but sharply transitioned out-of-phase outside the cluster. Specifically, during a Ca^2+^ influx event, Ca^2+^-triggered cAMP production dominated at the center of the AKAP79/150 cluster while Ca^2+^-triggered cAMP degradation was favored outside the cluster at the PM and in the center of the unit volume (**Fig. 4a**). Not surprisingly, the regime that recapitulates this phase relationship is sensitive to the spatially-restricted AC8/PDE1 balance and the diffusivity of cAMP. Assuming that AC8 clustering is driven by AKAP150:AC8 interactions, weakening this interaction would then reduce the AC8 cluster stabilization and lead to a redistribution of AC8 away from the nanoclusters and a decrease in the local concentration of AC8 within the clusters (**Fig. 4b**). Without the high local concentration of AC8 driving a net positive cAMP production within an AKAP79/150 cluster, the spatial domain at the PM where cAMP oscillates in-phase with Ca^2+^ is predicted to shrink while the out-of-phase regime expands and can reverse the phase at the cluster center (**Fig. 4c**).

**Figure 4.**
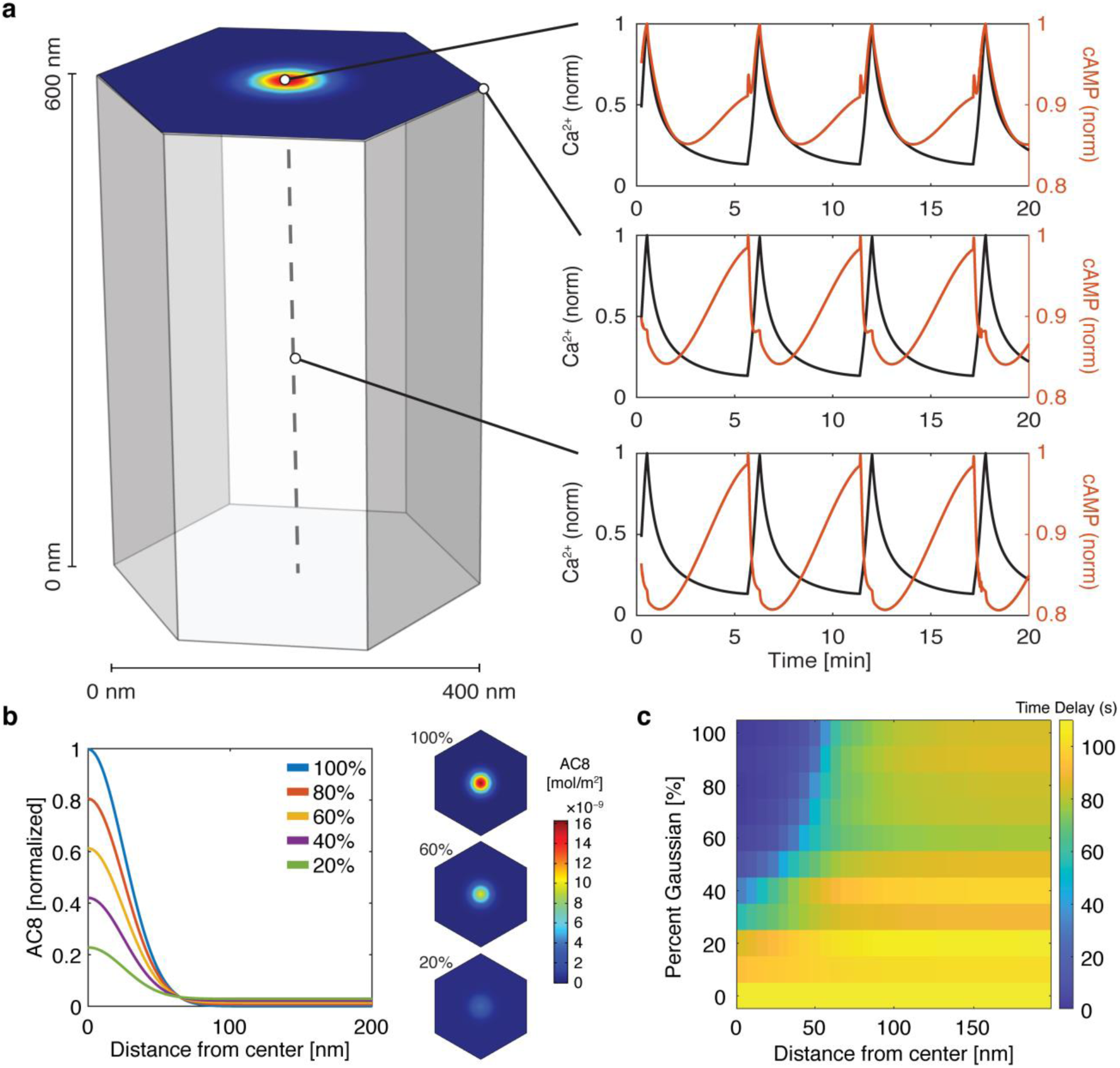
cAMP-Ca^2+^ phase relationship can be described by a 3D reaction-diffusion model involving clusters of AKAP79/150 and AC8. (a) 3D reaction-diffusion model with a single AKAP79/150:AC8 co-cluster positioned at the PM in the β cell in a hexagonal prism volume. cAMP oscillates in-phase immediately within the AKAP79/150:AC8 nanocluster due to the high effective concentration of AC8, but out-of-phase at the PM or cytosol due to the presence of PDE1 (cAMP – red, Ca^2+^ – blue). (b) Disruption of AKAP79/150:AC8 interaction can redistribute AC8 from within the cluster to the PM, shown by the half-Gaussian cross-sections and representative AC8 concentration heatmaps at the PM. (c) Heatmap depicting the time lag (s) for AC8 distribution (% Gaussian) and spatial distance (nm) from cluster center along PM.

To test this prediction, we overexpressed the amino terminus of AC8 (AC8^1-106^) required for interaction with AKAP79/150 (Willoughby et al. 2010) in order to compete with the binding of endogenous AC8 with the endogenous AKAP150 scaffold. The disruption of the AKAP150:AC8 interaction was validated by using proximity ligation assay (PLA) as an in situ assay for visualizing the interaction between AKAP150 and AC8. Compared to non-transfected cells, cells expressing the AC8^1-106^ peptide had a 39 ± 4% reduction in the number of PLA signals, indicating a decrease in AKAP150:AC8 interaction (**Extended Fig. 4**). Furthermore, STORM imaging showed that overexpression of the AC8^1-106^ peptide led to a decrease in the percentage of AC8 single molecule localizations within AC8 nanoclusters (n = 9) (**Fig. 5a**), consistent with the predicted redistribution of AC8 molecules (**Fig. 4b**). To test the impact of loss of AC8 molecules from the nanoclusters on the oscillation phase, we measured AKAP79/150-localized cAMP in the presence of AC8^1-106^ and observed a significant increase in the average lag time (43 ± 6sec, n = 33) (**Fig. 5b**). This is due to a higher proportion of cells exhibiting out-of-phase cAMP oscillations, indicating that the AKAP79/150:AC8 competitor peptide was sufficient in reversing the phase relationship in the AKAP79/150 compartment. Interestingly, we also observed many cells displaying irregular Ca^2+^ oscillations as indicated by a disruption in the periodic timing of individual cells’ Ca^2+^ peaks (**Fig. 5b**, left). This nanoscale perturbation establishes the regulatory role of the AKAP79/150:AC8 interaction in mediating the compartmentalized cAMP-Ca^2+^ phase relationship.

**Figure 5.**
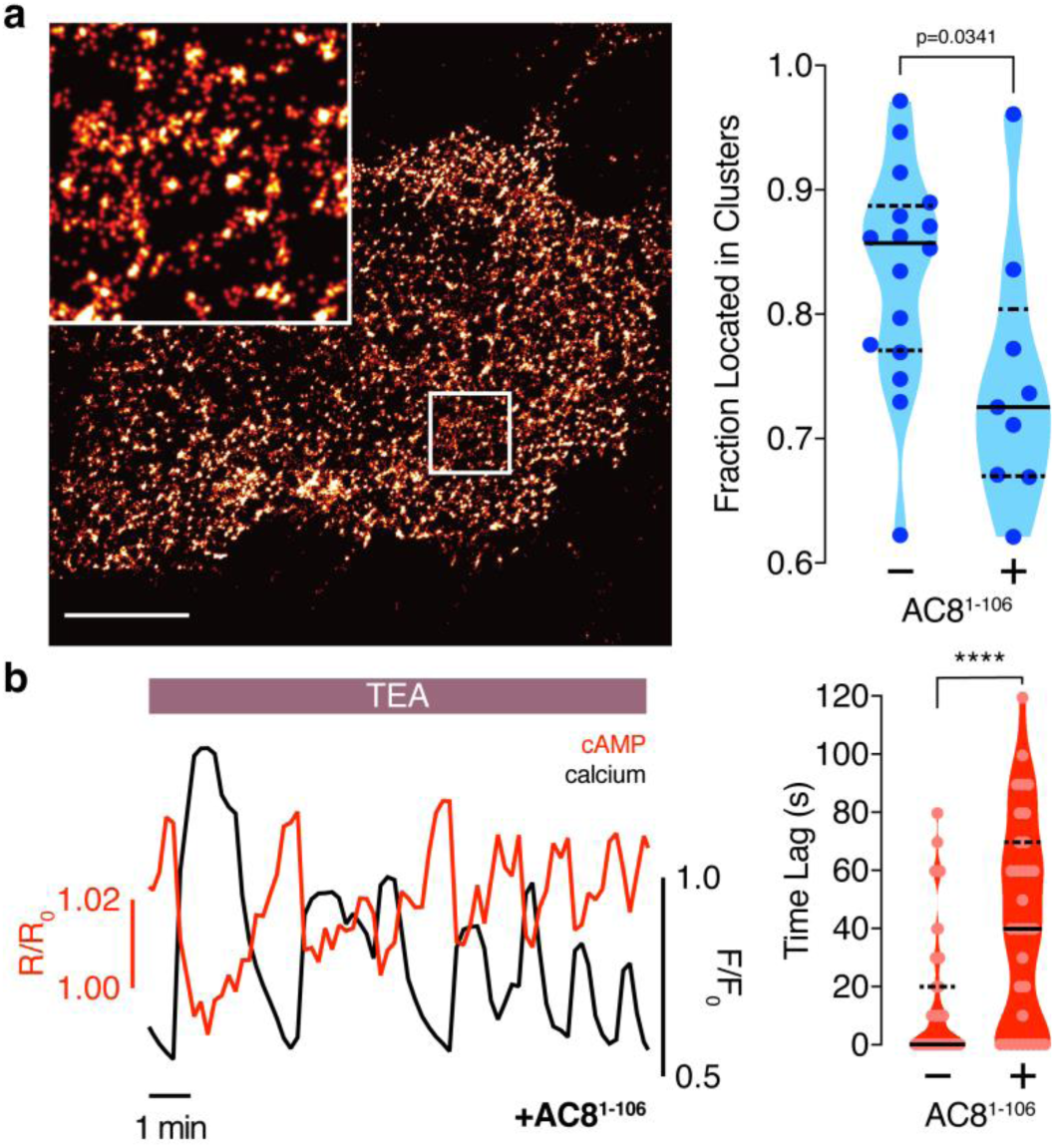
Disruption of the AKAP79/150:AC8 interaction is associated with a redistribution of AC8 at the PM and a phase shift of cAMP at the AKAP79/150 nanodomain. (a) Over-expression of the N-terminus of AC8 that is necessary and sufficient for mediating the AKAP79/150:AC8 interaction redistributes AC8 from within nanoclusters to the general PM, as seen in the STORM image (scale 5μm, inset 500nm) and measured by the percent of localizations that fall into nanoclusters. (b) Disruption of the AKAP79/150:AC8 interaction lengthens the time lag between the cAMP (red) and Ca^2+^ (black) signals at the AKAP79/150 compartment (avg. time lag in absence of disruptor is 13 ± 3 sec, n = 60, and presence of disruptor 43 ± 6 sec, n = 33, p<0.05) due to more cells displaying out-of-phase cAMP oscillations.

### AKAP79/150-mediated phase relationship is critical for regulating oscillatory Ca^2+^

Next we systematically examined the impact of perturbing the precisely regulated phase relationship within the AKPA79/150 compartment. Due to the modulatory role of PKA in the Ca^2+^-cAMP-PKA oscillatory circuit and the interaction between PKA and AKAP79/150, we wondered how the in-phase cAMP oscillations with respect to Ca^2+^ are translated into PKA activities and if spatial compartmentalization of the phase relationship is also maintained at the PKA activity level. Therefore, we extended our 3D model to include AKAP79/150-associated PKA (see Supplemental Information for model details). According to this extended model, PKA activity oscillations exhibit distinct phase relationships with respect to Ca^2+^ within and outside of the AKAP79/150 compartment (**Extended Fig. 5a**). To test this prediction, we fused our FRET-based biosensor for PKA activity (AKAR4) (Depry et al. 2011) to either full-length AKAP79 or the PM-targeting motif and expressed the sensors in MIN6 cells. Upon TEA stimulation, PKA activity was observed to oscillate with a lag time of time 25 ± 6sec (n = 15) within the AKAP79/150 compartment but with a lag time of 55 ± 8 sec (n = 12) (**Extended Fig. 5b-d**) at the general plasma membrane, indicating that the compartmentalized phase relationship is preserved from cAMP to PKA.

Spatiotemporal organization of PKA signaling and its phosphorylation targets via AKAPs have been implicated in regulating several important pathways. For example, PKA has been shown to phosphorylate Ca_V_1.2 in an AKAP79/150-dependent manner and this modification can influence the open probability of the channel (Murphy et al. 2014), suggesting a mechanistic link between local cAMP/PKA activity and global oscillatory Ca^2+^. Thus, we sought to study the functional role of the spatially-compartmentalized cAMP-Ca^2+^ phase relationship in regulating intracellular Ca^2+^ dynamics. We measured Ca^2+^ oscillations by RCaMP either in the presence of the EGFP-tagged AKAP79/150:AC8 disruptor peptide, AC8^1-106^, or EGFP alone as a control. Population-wide differences in Ca^2+^ dynamics, such as strength and timing, were observed in AC8^1-106^-transfected cells and visualized in heat maps depicting the normalized Ca^2+^ signal per cell versus time (**Fig. 6a**). Interestingly, we found that the expression of the disruptor peptide was correlated with a significant decrease in the peak ratio between the second Ca^2+^ peak and the first Ca^2+^ peak (control average -1.6%, n = 270; AC8^1-108^ average -10.8%, n = 562), post TEA addition, indicative of less sustained oscillations (**Fig. 6b,c**). In addition to intracellular Ca^2+^ concentration, the precise timing of internal oscillatory events is critical for modulating the β cell’s functions, such as glucose homeostasis and pulsatile insulin secretion (Fridlyand et al. 2010). In the presence of the disruptor peptide, cells also exhibited a longer elapsed time between oscillatory Ca^2+^ peaks (control average 3.9 ± 0.1 min, n = 270; AC8^1-108^ average 4.6 ± 0.1min, n = 562), suggesting that the timing of the signaling circuit was disturbed (**Fig. 6b,c**). In addition to the precise timing, the regularity of cytoplasmic Ca^2+^ in β cells is crucial in mediating pulsatile insulin secretion from the pancreas (Gilon et al. 2002; Schmitz et al. 2002). By stratifying the disruptor peptide-expressing cell population into “low, “medium,” and “high” expressers, and performing a blinded classification of responding cells based on the regularity of the Ca^2+^ oscillation (see Supplementary Information for details), we found a positive correlation between the percentage of cells exhibiting irregular oscillations and the expression level of the disruptor peptide (42% for low-expressing vs. 68% for high-expressing AC8^1-106^ disruptor) (**Fig. 6d**). Taken together, these data signify that the compartmentalized cAMP-Ca^2+^ phase relationship regulates the oscillatory Ca^2+^ signal and plays an important role in determining the pace, regularity, and sustainability of the Ca^2+^ oscillations.

**Figure 6.**
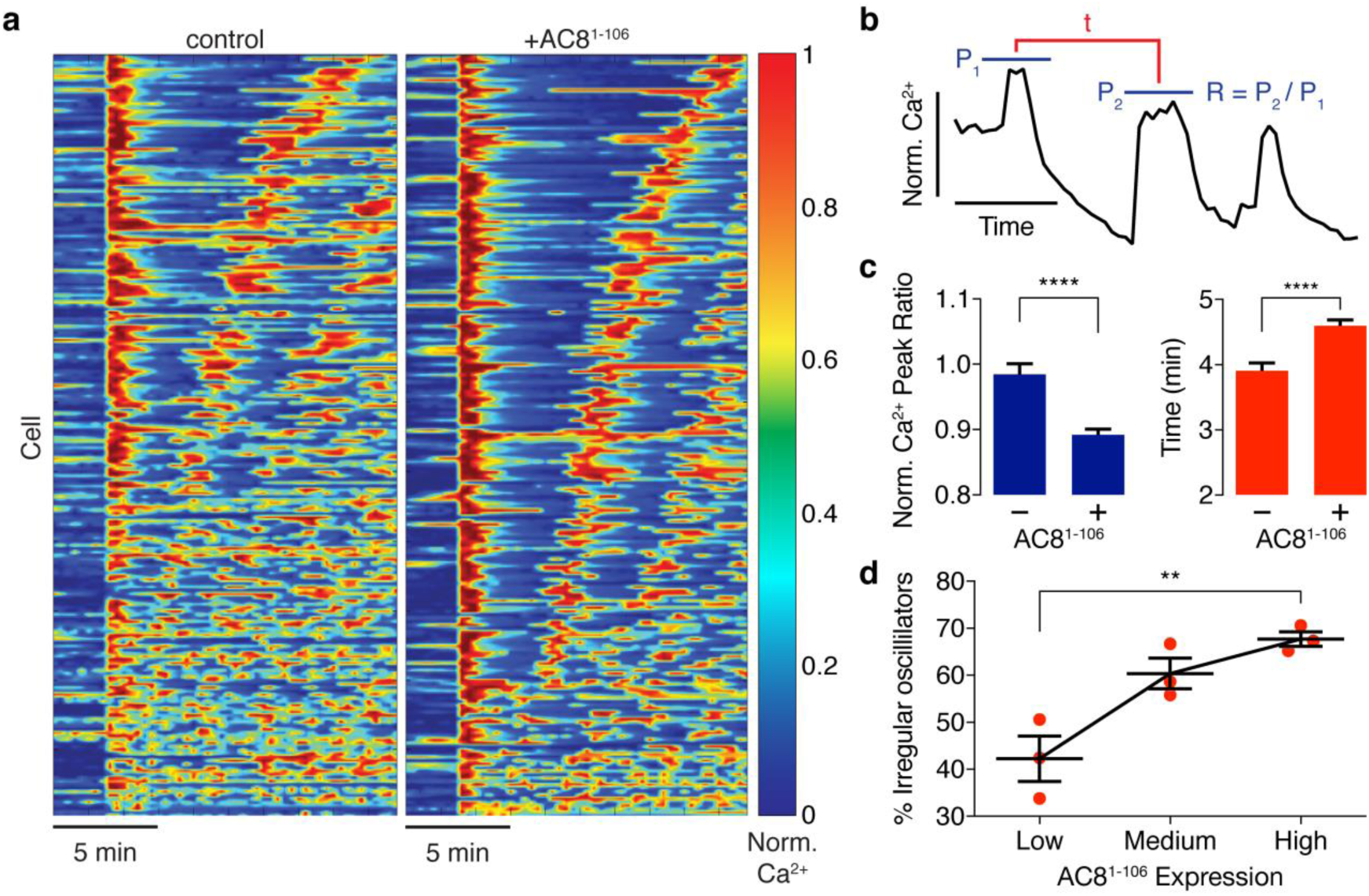
Ca^2+^ oscillatory dynamics are affected by expression of the disruptor peptide in β cells. (a) Heatmap depicting Ca^2+^ oscillations for 220 randomly selected cells with EGFP alone co-expressed (control) or EGFP-tagged AC8^1-106^ (AKAP79/150:AC8 disruptor), ordered by a mixed parameter describing the time lag between the first two Ca^2+^ peaks and the avg. timelag between all Ca^2+^ peaks. (b) Schematic describing two Ca^2+^ oscillatory parameters: the ratio between the first two Ca^2+^ peaks (R = P_2_ / P_1_) and the interpeak timing (t). (c) The peak ratio is decreased in the presence of the AKAP79/150:AC8 disruptor, indicating less of a sustained Ca^2+^ oscillatory response. Over-expression of the disruptor also lengthens the timing between peaks. (d) Expression level of the disruptor is correlated with an increase in the percentage of cells eliciting irregular oscillations (total n = 562).

## DISCUSSION

Biological oscillations represent a rich way of encoding information. Here we show the phase in an oscillatory signaling circuit, like the amplitude and frequency, represents a novel mode of informational encoding which itself can be spatiotemporally regulated. In the case of the Ca^2+^-cAMP-PKA circuit, the oscillatory cAMP/PKA phase relative to a widespread Ca^2+^ signal is distinctly regulated within two adjacent membrane compartments through intracellular organization of scaffolds and signaling effectors. Localized perturbation of this spatial phase signature disrupts global Ca^2+^ oscillations and thus has far-reaching consequences on the functional landscape of the β cell.

Compartmentalization of cAMP/PKA signaling is instrumental in processing a diverse set of inputs and mediating specific cellular functions; however, the mechanistic details of compartmentalization are still largely unresolved (Mesheshe et al. 2018). Given the measured kinetic rates of most ACs and PDEs, coupled with apparent fast diffusion of the small cAMP molecule, the generation of local cAMP gradients around single enzymes is unfeasible (Conti et al. 2014). Context-dependent discrepancies in some of the kinetics (i.e. differences of *in vitro* versus *in vivo* measurements) or slower cAMP diffusion due to buffering have been proposed as potential mechanisms for cAMP compartmentalization (Agarwal et al. 2016). Here we propose that the nanoscale organization of key cAMP effectors and regulators as a novel mechanism for cAMP compartmentation. Despite the slow rates measured for individual ACs, we computationally and experimentally describe conditions in which the generation of compartmentalized cAMP can emerge from the clustering of many AC8 enzymes at the membrane and bulk distribution of PDE1 in the cytoplasm. Alternations to this nanoscale organization lead to dysregulated Ca^2+^ oscillations, demonstrating the functional importance in maintaining this organization. This system also serves as a nanoscale demonstration of how a cell can translate a global signal (Ca^2+^) into a compartmentalized signal (cAMP/PKA activity) by local activation and global inhibition, a strategy that is likely utilized in many other cellular contexts (Levchenko et al. 2000; Purvis et al. 2014).

Multiple mechanisms could contribute to the signaling and functional effects controlled by the compartmentalized cAMP-Ca^2+^ phase mediated by AKAP79/150 in β cells. AKAPs can recruit PKA to regulate channel activities (Dell’Acqua et al. 2006; Mo et al. 2017; Torres-Quesada et al. 2017), such as in the regulation of voltage-mediated Ca^2+^ entry via PKA-dependent phosphorylation of Ca_V_1.2 (Murphy et al. 2014) or the modulation of store-operated Ca^2+^ entry by both PKA-dependent STIM1 and Orai1 phosphorylation (Thompson et al. 2015; Zhang et al. 2019). Additional levels of regulatory feedback within the Ca^2+^-cAMP-PKA oscillatory circuit have also been identified, such as a negative feedback loop involving PKA phosphorylation of AC8, thereby fine-tuning the circuit dynamics (Willoughby et al. 2012). Localized cAMP/PKA signaling at the AKAP79/150 scaffold might also play a role in directly regulating downstream insulin secretion due to close interactions between AKAP79/150 and the insulin secretory granules via Ca_V_1.2 (Barg et al. 2001). Several important processes and components of the secretory machinery have been identified as targets of PKA signaling here, such as PKA-dependent mobilization of granules (Renstrom et al. 2004) and modulation of the synaptosomal protein SNAP25 (Gao et al. 2016). In addition to PKA-dependent secretory control, cAMP has recently been implicated to play a role in fusion pore formation via a cAMP-regulated guanine exchange factor Epac (Gucek et al. 2018). Compartmentalized cAMP/PKA signaling at the AKAP79/150 macromolecular complex is likely involved in the regulation of many β cell processes and more work will be needed to further establish the link between the oscillatory circuit and the mechanisms involved in decoding the information embedded in the local phase relationship.

The Ca^2+^-cAMP-PKA oscillatory circuit in pancreatic β cells integrates many important regulators of cellular function, and the precise coordination of each is required for proper signaling control. Here we have uncovered a spatiotemporal organization of the circuit where the oscillatory phase between cAMP/PKA and Ca^2+^ depend on the spatial proximity of the AKAP79/150 scaffold protein and AC8. The construction principles of this signaling nanodomain, including the spatial distributions of sinks and sources, likely represent a generalized strategy for the generation of other compartmentalized signals and provide a unique modality in which cells embed, process, and produce signaling information.

## Acknowledgements

We thank Dr. John D. Scott for providing AKAP79, Dr. Dermot Cooper for providing (Ci/Ce)Epac2-camps and AC8, Dr. Loren Looger for providing RCaMP and Dr. Jochen Lang for providing the shAC8 construct. We thank Dr. Susan Taylor for critical reading of the manuscript. This work was supported by R01 DK073368 (to J. Z.), DOD AFOSR FA9550-18-1-0051 (to P.R. and J. Z.) and ONR N00014-17-1-2628 (to P.R.).

## Author Contributions

B.T., S.M., and J. Z. conceived of the study. B.T. and J.Z. designed the experiments. B.T. performed all of the biosensor imaging and biochemical experiments. M.G., D.O. and P.R. built the computational models. B.T. and B.R. performed the STORM imaging, and C.H.B., B.T., and E.G. analyzed the STORM imaging data. J.Z., P.R., and J.X. supervised the research. B.T. and J.Z. wrote the manuscript, with input from all authors.

## Materials and Methods

### Gene Construction

For AKAP79-(Ci/Ce)Epac2-camps, AKAP79 (from Dr. John D. Scott) was PCR amplified to have HindIII/BamHI digestion sites and (Ci/Ce)Epac2-camps (from Dr. Dermot Cooper) was PCR amplified to have BamHI/EcoRI digestions sites. Both fragments were inserted into pcDNA3 (Invitrogen) backbone for mammalian expression (cAMP sensor is C terminal to AKAP79). For AKAP79-AKAR4, a similar approach was taken where AKAR4 was dropped between BamHI/EcoRI. For AC8 (from Dr. D. Cooper), AC8^1-108^. Gibson Assembly was used to insert the genes into the pcDNA3 mammalian expression vector. The shAC8 construct for AC8 knockdown was previously verified and a gift from Dr. Jochen Lang. RCaMP was a gift from Dr. Loren Looger.

### Cell Culture

MIN6 cells (a mouse insulinoma β cell line) were plated onto sterilized glass coverslips in 35-mm dishes and grown to 50–90% confluency in DMEM (10% FBS, 4.5g/L glucose) at 37°C with 5% CO2. Cells were transfected using Lipofectamine 2000 (Invitrogen) for 20–48 h before imaging.

### Imaging

Cells were washed twice with Hanks’ balanced salt solution buffer and maintained in the dark at room temperature. Cells were imaged on a Zeiss Axiovert 200M microscope with a cooled charge-coupled device camera (MicroMAX BFT512, Roper Scientific, Trenton, NJ) controlled by METAFLUOR 6.2 software (Universal Imaging, Downingtown, PA). Dual cyan/yellow emission ratio imaging used a 420DF20 excitation filter, a 450DRLP dichroic mirror, and two emission filters [475DF40 for CFP and 535DF25 for YFP]. RFP fluorescence was imaged using a 568D55 excitation filter, a 600DRLP dichroic mirror, and a 650DF100 emission filter. GFP fluorescence was imaged using a 480DF30 excitation filter, a 505DRLP dichroic mirror, and a 535DF45 emission filter. These filters were alternated by a filter-changer Lambda 10–2(Sutter Instruments, Novato, CA). Exposure time was 50–500 ms, and images were taken every 10–30 s. Fluorescence images were background-corrected by subtracting the fluorescence intensity of background with no cells from the emission intensities of cells expressing fluorescent reporters. The ratio of yellow/cyan emission, RFP intensity, and GFP intensity were then calculated at different time points. The values of all time courses were normalized by dividing each by the average basal value before drug addition. Custom Java code, MATLAB scripts, and CellProfiler (Broad Institute) pipelines were written to segment cells, select ROIs, and analyze traces.

For confocal imaging, images were collected with a C2 plus on a Nikon Ti2 inverted microscope equipped with a Plan Apo lambda 60x oil immersion objective NA 1.4. YFP fluorescence fluorescence was excited with the 488nm line from a LU-N4 laser. Images were acquired with a DUVB detector collecting emission from 495nm to 600nm with a virtual spectral GaAsP detector controlled by NIS Elements software. The pinhole was set at 30μm. Frame size was 1024 x 1024pix.

### Super-resolution Imaging (STORM)

For fixed-cell stochastic optical reconstruction microscopy (STORM) imaging, cells were fixed with 4% paraformaldehyde (PFA) and 0.2% glutaraldehyde (GA) for 20 min and then washed with 100 mM glycine in Hanks’ balanced salt solution (HBSS) to quench the free PFA. Cells were permeabilized and blocked in a permeabilization solution with 0.1% Triton X-100, 0.2% bovine serum albumin, 5% goat serum, and 0.01% sodium azide in HBSS. The cells were then incubated overnight at 4°C with an anti-AC8 antibody (Abcam, ab196686) at a 1:2000 dilution or an anti-AKAP150 (Millipore Sigma, 07-210) antibody at a 1:500 dilution, followed by 1 to 2 hours with goat anti-rabbit Alexa 647–conjugated antibody (ThermoFisher Scientific, A21245) at 1:1000 dilution. The cells were then post-fixed again in 4% PFA and 0.2% GA, quenched with 100 mM glycine in HBSS, and washed with HBSS to prepare for imaging. Immediately before imaging, the medium was changed to STORM-compatible buffer (50 mM tris-HCl (pH 8.0), 10 mM NaCl, and 10% glucose) with glucose oxidase (560 μg/ml), catalase (170 μg/ml), and mercapto-ethylamide (7.7 mg/ml). STORM images were obtained using a Nikon Ti total internal reflection fluorescence (TIRF) microscope with N-STORM, an Andor IXON3 Ultra DU897 EMCCD, and a 100× oil immersion TIRF objective. Photoactivation was driven by a Coherent 405-nm laser, while excitation was driven with a Coherent 647-nm laser. All image analysis and image reconstruction were performed using both Nikon Elements analysis software and custom-written MATLAB scripts. Blinking correction was performed by implementing the pairwise Distance Distribution Correction (DDC) algorithm (Bohrer et al. 2019). Cluster property measurements were performed using Ripley-K analysis and custom mean-shift code for segmentation.

### Proximity Ligation Assay

Antibodies for AC8 and AKAP150, mentioned in STORM section, were buffer exchanged into DPBS and conjugated with MINUS or PLUS oligos, following the Sigma DuoLink in situ Probemaker kits. PLA experiments were performed using the Duolink® in situ red kit for proximity ligation assays according to the provided protocol. The only protocol modification was to extend the amplification time by 50 min. Briefly, cells were fixed and permeabilized as in the STORM experiments before incubation with PLUS and MINUS oligo-conjugated primary antibodies for 30 min at 37°C each with washes after each step. Ligation of the nucleotides and amplification of the strand occurred sequentially by incubating cells with first ligase then polymerase and detection solution. PLA experiments with AKAP95 antibodies from different species were used as positive controls in HEK293T cells, and experiments with just one oligo-labeled primary antibody or the other were our negative control. Images were acquired on a Nikon Ti Eclipse epifluorescence scope with z-control and maximum intensity projections were created. A cross section of the nucleus (3.6-5 μm) was also acquired and the number of dots per cell was counted using the nucleus as reference.

### Computational Modeling

See Supplementary Information.

## Supplemental Information

### cAMP Analysis and Quantification

AKAP79-(Ci/Ce)Epac2-camps transfected cells displayed cAMP oscillations that were either in-phase or out-of-phase with their respective Ca^2+^ signal. This was in sharp contrast to responsive lyn-(Ci/Ce)Epac2-camps transfected cells where all cells yielded only out-of-phase oscillations. We found a strong correlation between the AKAP79-fused sensor expression level and the observed cAMP-Ca^2+^ phase relationship, with cells having lower levels of sensor present displaying predominantly in-phase cAMP oscillations and cells with higher levels of the AKAP79/150-fused biosensor exhibiting out-of-phase oscillations **(Accessory Fig. 1)**.

Overexpression of the AKAP79 scaffold likely changed the stoichiometry of the signaling complexes and resulted in unsuccessful targeting of the biosensor to functional AKAP79/150 domains, and so in this manuscript we considered only TEA-responsive cells below an AKAP79 expression threshold determined by the YFP acceptor fluorescence (**Accessory Fig. 1**).

### Time Lag Calculation

Due to the heterogeneity of cellular Ca^2+^ and cAMP/PKA activity oscillatory responses in each cell (i.e. variations in frequency, amplitude, and regularity), we sought an applicable metric to describe the phase relationship. Here we measure the lag time (sec) between the Ca^2+^ signal trace and the cAMP/PKA activity signal trace. Specifically, we high-pass filtered both the Ca^2+^ and cAMP/PKA activity traces (approx.. 20 min) to subtract out slowly varying baseline changes, normalized the traces so that the maximum intensity/FRET ratio was set to 1, and then computed the cross-correlation to measure the signal overlap for different lag times. To calculate the lag time, we identified peaks in the cross-correlation passing a peak prominence cutoff and found the absolute value of the shortest lag time corresponding to a peak maximum. For in-phase oscillations, the lag time was typically small (τ ≤ 20 sec) due to the two signal traces oscillating in synchrony. However, out-of-phase oscillations typically corresponded to longer lag times (τ > 20 sec) due to the anti-phasic relationship seen in the peak timing and peak shape. Analysis was performed with custom scripts in MATLAB and Java, and pipelines in CellProfiler.

### Quantification of Nanodomain Perturbation Effects on Global Ca^2+^

In order to measure the effects of AKAP79/150:AC8 disruption on Ca^2+^ dynamics, we transiently transfected and expressed AC8^1-106^ in MIN6 and measured Ca^2+^ with RCaMP. For quantification of the interpeak timing and peak ratio, we first selected cells that responded to the TEA treatment, identified Ca^2+^ peaks passing a peak prominence cutoff, and finally calculated the avg. time between peak maxima and RFP intensity ratio between the second and first Ca^2+^ peak maxima. To find the percentage of cells with regular vs. irregular Ca^2+^ oscillations, we randomized all Ca^2+^ traces from the experimental and control samples and performed a blinded classification to sort the single cell traces as regular, irregular, or nonresponsive. Analysis was performed with custom scripts in MATLAB and Java, and pipelines in CellProfiler.

### Computational Modeling

#### Well-mixed system

#### Assumptions

- Signaling components are present in large enough quantities that concentration changes are smooth and move in a deterministic fashion.
- Well-mixed kinetic rate constants contain conversion factors between compartments, i.e membrane to cytosol.
- Binding interactions occur rapidly enough such that any kinetic parameter, *k*, remains constant on surfaces (Berry 2002).
- A-kinase-anchoring protein (AKAP) does not alter the activity of the catalytic subunits of Protein kinase A (PKA) instead it only affects localization.
- Ca^2+^ independent activity of Adenylyl cyclase (AC) is of the same strength as inactive Ca^2+^ dependent Adenylyl cyclase (AC8) and stays at a constant value.

### Modeling well-mixed chemical reactions

#### Mass-action kinetics

We generated an ordinary differential equation (ODE) for every species using mass-action kinetics for each binding interaction. The law of mass action states that the rate of a chemical reaction is proportional to the product of the concentration of the reactants raised to the power of their stoichiometric coefficient (Guldberg and Waage 1879). Mass action kinetics rely on the assumption that the rate constant, *k*, is constant over time. For example, consider the one-reaction system:

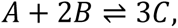

where the forward and backward rates are *k*_1_ and *k*_2_. The differential equations describing the dynamics of species *A*, *B*, and *C* under mass-action kinetics are:

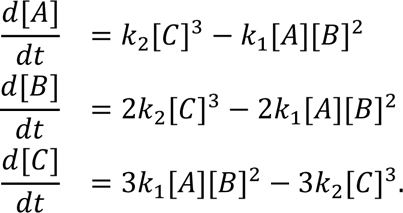

#### Michaelis-Menten kinetics

We used Michaelis-Menten kinetics to model the kinetics of enzyme-catalyzed reactions. When a reaction is catalyzed by an enzyme with kinetic properties *k_cat_* and *K_M_*,

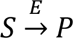

then the reaction rate is given by

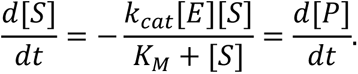

For Michaelis-Menten kinetics, concentrations of reactants and products must be in large enough quantities, and one of the following conditions must apply: the concentration of the substrate is much larger than the concentration of products, [S]≫[P], and/or the energy released in the reaction is very large, *ΔG* ≪ 0.

#### Model Development

We constructed a biochemical network to represent interactions between Ca^2+^ and cAMP in *β*-cells, **Figure 1B**, (Ni et al. 2011; Fridlyand and Philipson 2016) after a depolarization event. The computational model considered the dynamics of calcium, potassium, leaky, and calcium-sensitive potassium channels (**Table S1**). Importantly, we included feedback of PKA with KATP channels and the inclusion of Ca^2+^-sensitive ACs and PDEs. The model contains **92** parameters with **11** free parameters. The values of the parameters are constrained through both previously peer-reviewed publication results (Ni et al. 2011; Boras et al. 2014; Lai et al. 2015; Masada et al. 2009; Ang and Antoni 2002) and with new experimental results using obtained FRET measurements. To constrain source and sink activation rates, previously published literature of AC and PDE stimulus-response curves (of related isoforms) was utilized (Masada et al. 2009; Ang and Antoni 2002). COPASI was used to calculate initial guesses for kinetic activation parameters. FRET measurements took precedence over binding curves, especially for the spatial model (section 3) where further fitting routines were performed to refine the model. Model variations were performed to attain both semi-physiological concentrations (within the ranges of the sensor) and phase (period and relation) information. Predictions are made on qualitative behavior as opposed to quantitative as proper parameter fitting would require much more data than available for this system.

Values and reaction sets used in the well-mixed model can be found in **Tables S1-S5**. The network of interactions was constructed using COPASI (version 4.23, build 184) (http://www.nrcam.uchc.edu, http://copasi.org/). The model was built in COPASI to leverage the inbuilt fitting techniques for initial parameter guesses pre-FRET.

#### Well-mixed computational results

The network shown in **Figure 1B** has been shown to exhibit oscillations through cAMP variation due to the action of PKA on IP3 receptors and KATP Plasma membrane channels. This network has been studied in many labs previous work (Ni et al. 2011; Fridlyand, Tamarina, and Philipson 2003; Fridlyand and Philipson 2011, 2016; Han et al. 1999) and has been explored to show:

- The system can be moved in and out of phase through tuning of AC (Fridlyand and Philipson 2016).
- The oscillation rate can be tuned through PKA feedback to KATP channels (Ni et al. 2011).
- Both Ca^2+^ sensitive PDE and AC is necessary for oscillations to occur (Ni et al. 2011).

Our model results agree with the above findings.

Variations in the connection strength of sources and sinks also introduced another finding– changing the component’s (source or sink) variability in the regime of Ca^2+^ spiking (0.1-1.2*μM*) a switch in phase can also be observed, **Accessory Fig. 2**. Compared to well-mixed results, **Accessory Fig. 2a**, by decreasing the activation rate of Ca_2_AC from 56 to 0.6 *s*^−1^, a switch in phase can occur, **Accessory Fig. 2c**. Yet, we notice how increasing PDE activity does not switch the phase **Accessory Fig. 2b**, only after subsequently decreasing PDE and CaM association does cause the phase change **Accessory Fig. 2d**. This is due to both changes being required for the activity variation of PDE to outperform that of AC. These findings lead to more questions about how source and sink activities relate in a variable Ca^2+^ regime.

#### Well-mixed reaction tables

**Table S1:** Voltage Gated Channel Reactions

| # | Species | Expression | Parameters | Ref. |
| --- | --- | --- | --- | --- |
| 1 | Membrane Voltage | $\frac{dV}{dt} = \frac{-I_{Ca} - I_K - I_L - I_{KCa}}{C_m}$ | $C_m=5.3$ pF | (Ni et al. 2011) |
| 2 | Ca<sup>2+</sup> current | $I_{Ca} = g_{Ca}m_{\infty}(V - E_{Ca})$ | $g_{Ca}=600$ pS,<br>$E_{Ca}=100$ mV | (Ni et al. 2011) |
| 3 | Fraction of open VGCC<br>(at steady state) | $m_{\infty} = \frac{1}{2} \left( 1 + \tanh \left( \frac{V - v_1}{v_2} \right) \right)$ | $v_1=-20$ mV, $v_2=24$<br>mV | (Ni et al. 2011) |
| 4 | K<sup>+</sup> current | $I_K = g_K w (V - E_K)$ | $g_K=240$ pS, $E_{Ca}=-$<br>75 mV | (Ni et al. 2011) |
| 5 | Fraction of open K<sup>+</sup><br>channels (at steady<br>state) | $\frac{dw}{dt} = \frac{\phi(w_{\infty} - w)}{\tau}$ | $\phi=35 \frac{1}{s}$ | (Ni et al. 2011) |
| 6 | Time constant for K<sup>+</sup><br>channel open<br>probability | $\tau = \frac{1}{\cosh \left( \frac{V - v_3}{2v_4} \right)}$ | $v_3=-16$ mV<br>$v_4=11.2$ mV | (Ni et al. 2011) |
| 7 | leak current | $I_L = g_L(V - E_L)$ | $g_L=150$ pS, $E_L=-$<br>75 mV | (Zhang et al.<br>2005) |
| 8 | Ca<sup>2+</sup> gated K<sup>+</sup> current | $I_{KCa} = g_{KCa} \frac{Ca}{Ca + K_{KCa}} (V - E_K)$ | $g_{KCa}=2000$ pS,<br>$E_{Ca}=-75$ mV,<br>$K_{KCa}=5$ $\mu M$ | (Ni et al. 2011) |

**Table S2:**
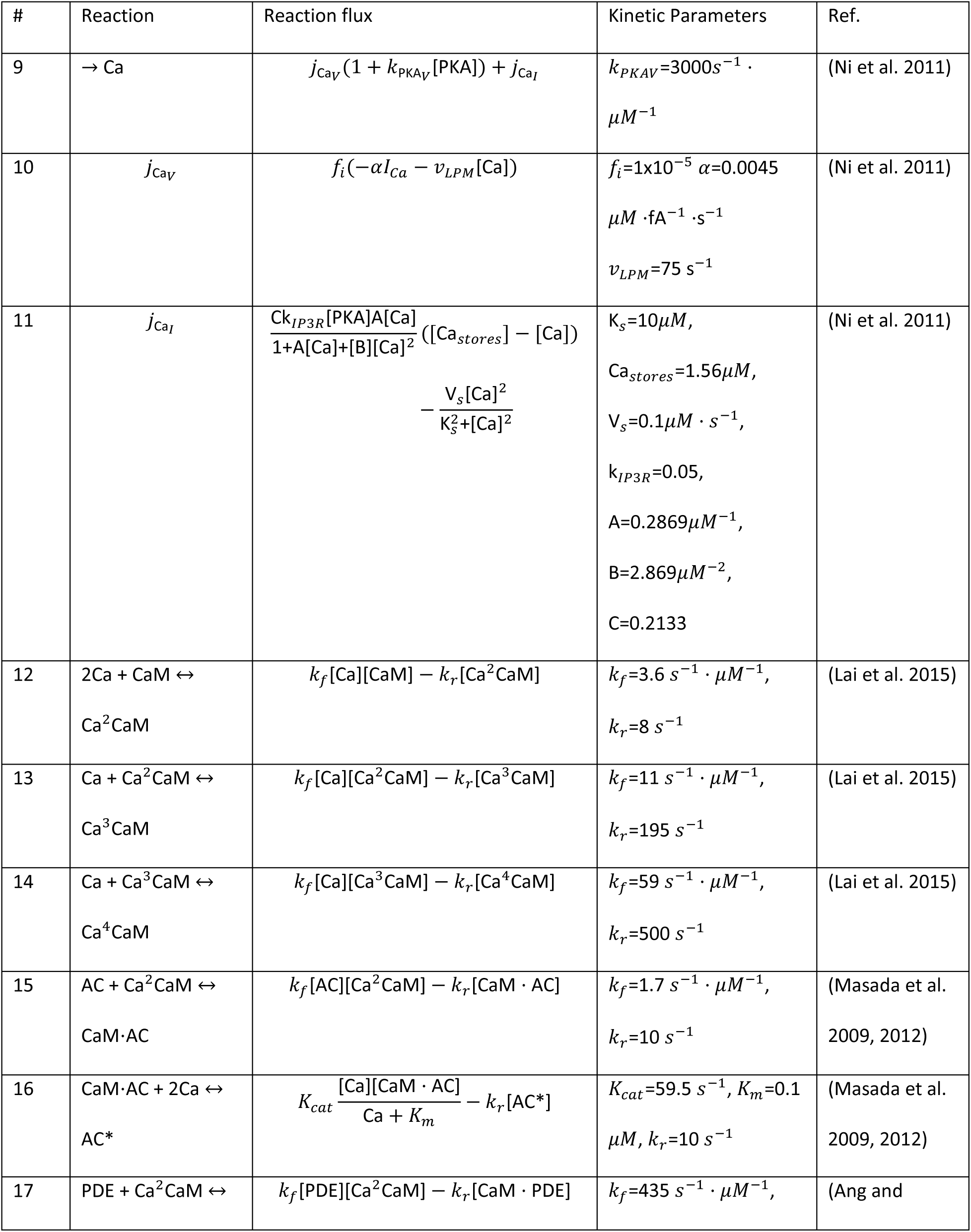
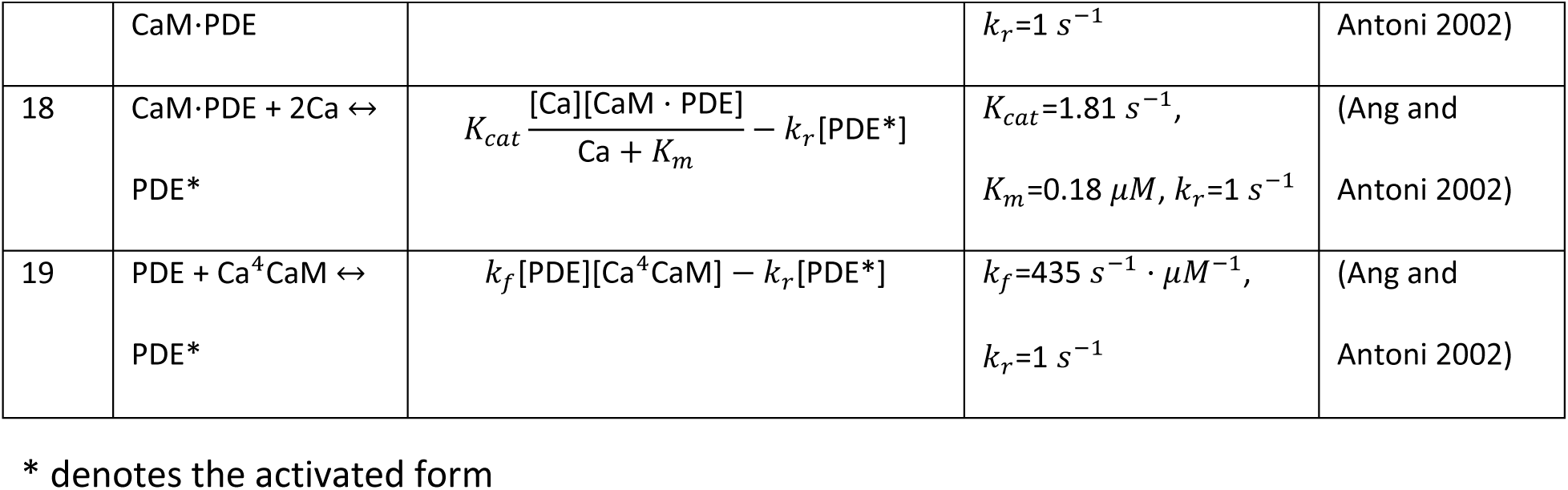
Ca^2+^ Flux and Reactions

**Table S3:** cAMP Reactions

| # | Reaction | Reaction Flux | Kinetic Parameters | Ref. |
| --- | --- | --- | --- | --- |
| 20 | $\rightarrow \text{cAMP}$ | $k_{base}([CaM \cdot AC] + [AC] + [AC_{ind}]) + k_{act}[AC^*]$ | $k_{base}=0.1 \text{ s}^{-1},$<br>$k_{act}=0.785 \text{ s}^{-1}$ | (Ni et al. 2011; Fridlyand and Philipson 2016) |
| 21 | cAMP→ | $k_{base}[cAMP] \frac{[CaM \cdot PDE] + [PDE]}{[cAMP] + K_m} + k_{act} \frac{[cAMP][PDE^*]}{[cAMP] + K_m}$ | $k_{base}=0.2 \text{ s}^{-1},$<br>$K_m = 0.6 \mu M$<br>$k_{act}=2.5 \text{ s}^{-1}$ | (Ni et al. 2011; Fridlyand and Philipson 2016) |
| 22 | cAMP→ | $V_{ind} \frac{[cAMP]}{[cAMP] + K_m}$ | $V_{ind}=2.5 \mu M \cdot \text{s}^{-1},$<br>$K_m = 1.4 \mu M$ | (Ni et al. 2011; Fridlyand and Philipson 2016) |
| 23 | cAMP + R2 → R2<sub>b</sub> | $k_f[cAMP][R2] - k_r[R2_b]$ | $k_f=1 \text{ s}^{-1} \cdot \mu M^{-1},$<br>$k_r=0.00033 \text{ s}^{-1}$ | (Boras et al. 2014) |
| 24 | cAMP + R2<sub>b</sub> → R2<sub>ba</sub> | $k_f[cAMP][R2_b] - k_r[R2_{ba}]$ | $k_f=1 \text{ s}^{-1} \cdot \mu M^{-1},$<br>$k_r=0.00105 \text{ s}^{-1}$ | (Boras et al. 2014) |
| 25 | $\text{cAMP} + \text{R2}_b \rightarrow \text{R2}_{bb}$ | $k_f[\text{cAMP}][\text{R2}_b] - k_r[\text{R2}_{bb}]$ | $k_f=1 \text{ s}^{-1} \cdot \mu\text{M}^{-1},$<br>$k_r=0.00132 \text{ s}^{-1}$ | (Boras et al. 2014) |
| 26 | $\text{cAMP} + \text{R2}_{ba} \rightarrow \text{R2}_{bba}$ | $k_f[\text{cAMP}][\text{R2}_{ba}] - k_r[\text{R2}_{bba}]$ | $k_f=1 \text{ s}^{-1} \cdot \mu\text{M}^{-1},$<br>$k_r=0.0013 \text{ s}^{-1}$ | (Boras et al. 2014) |
| 27 | $\text{cAMP} + \text{R2}_{bb} \rightarrow \text{R2}_{bba}$ | $k_f[\text{cAMP}][\text{R2}_{bb}] - k_r[\text{R2}_{bba}]$ | $k_f=1 \text{ s}^{-1} \cdot \mu\text{M}^{-1},$<br>$k_r=0.00103 \text{ s}^{-1}$ | (Boras et al. 2014) |
| 28 | $\text{cAMP} + \text{R2}_{bba} \rightarrow \text{R2}_{bbaa}$ | $k_f[\text{cAMP}][\text{R2}_{bba}] - k_r[\text{R2}_{bbaa}]$ | $k_f=1 \text{ s}^{-1} \cdot \mu\text{M}^{-1},$<br>$k_r=0.0114 \text{ s}^{-1}$ | (Boras et al. 2014) |
| 29 | $\text{PKA} + \text{R2} \rightarrow \text{R2C}$ | $k_f[\text{PKA}][\text{R2}] - k_r[\text{R2C}]$ | $k_f=1 \text{ s}^{-1} \cdot \mu\text{M}^{-1},$<br>$k_r=1.26\text{E-}7 \text{ s}^{-1}$ | (Boras et al. 2014) |
| 30 | $\text{PKA} + \text{R2}_b \rightarrow \text{R2}_b\text{C}$ | $k_f[\text{PKA}][\text{R2}_b] - k_r[\text{R2}_b\text{C}]$ | $k_f=1 \text{ s}^{-1} \cdot \mu\text{M}^{-1},$<br>$k_r=2.52\text{E-}7 \text{ s}^{-1}$ | (Boras et al. 2014) |

**Table S4:**
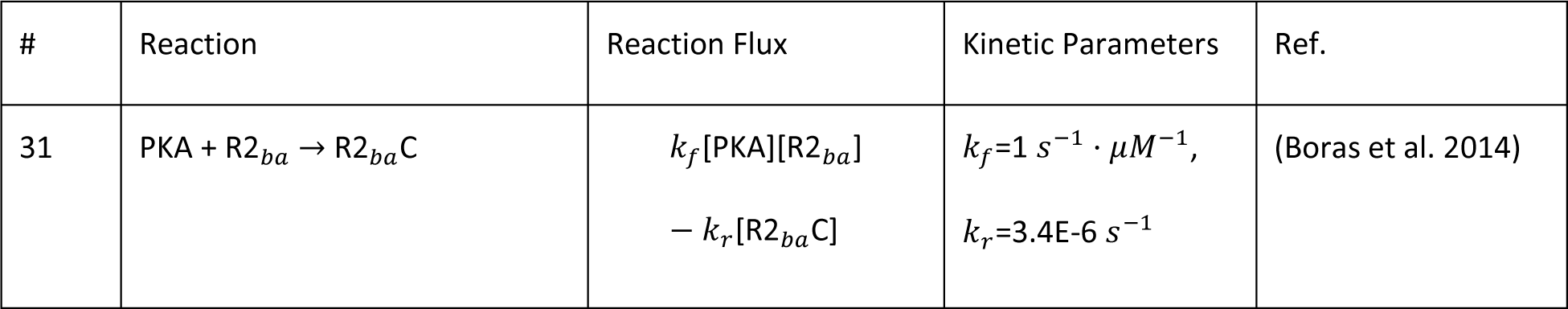
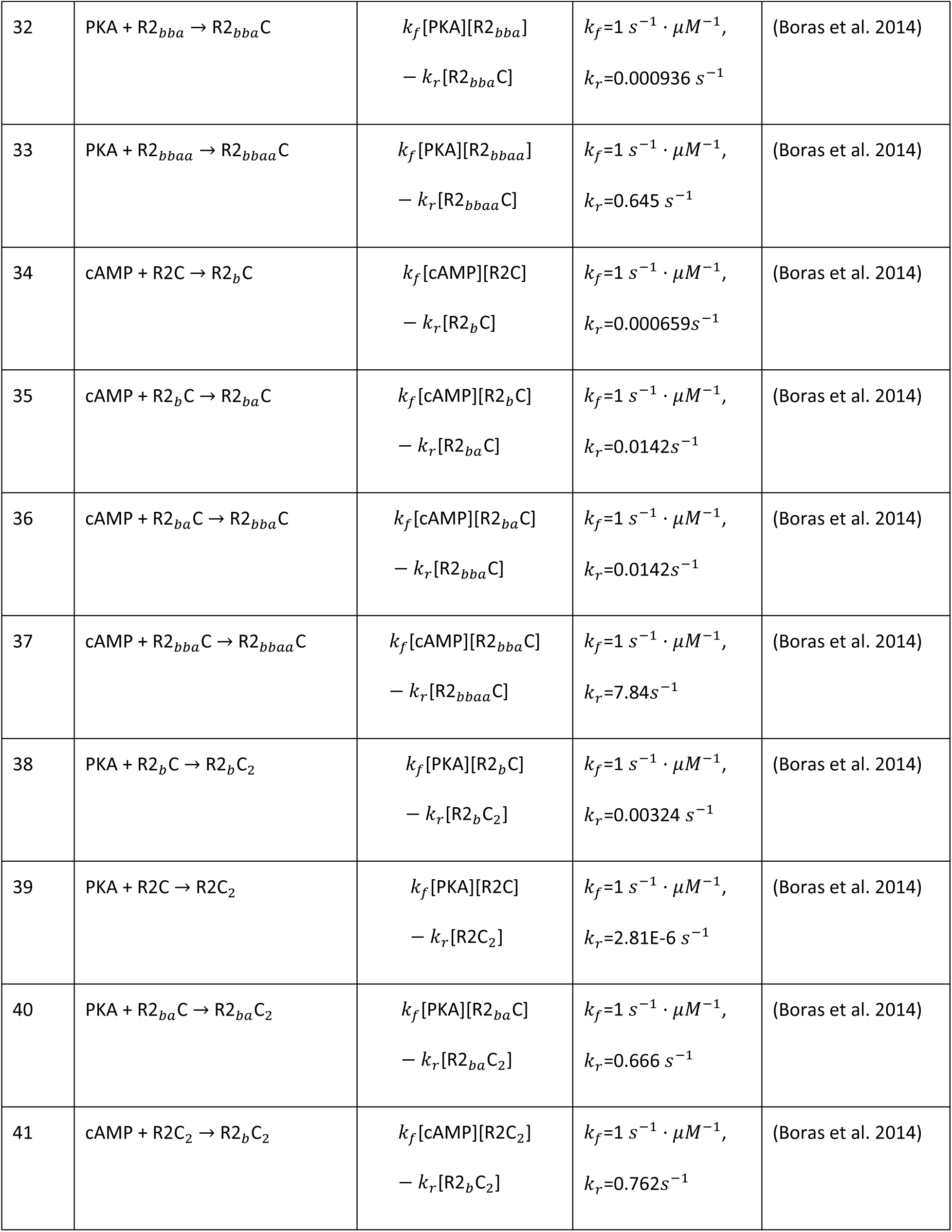
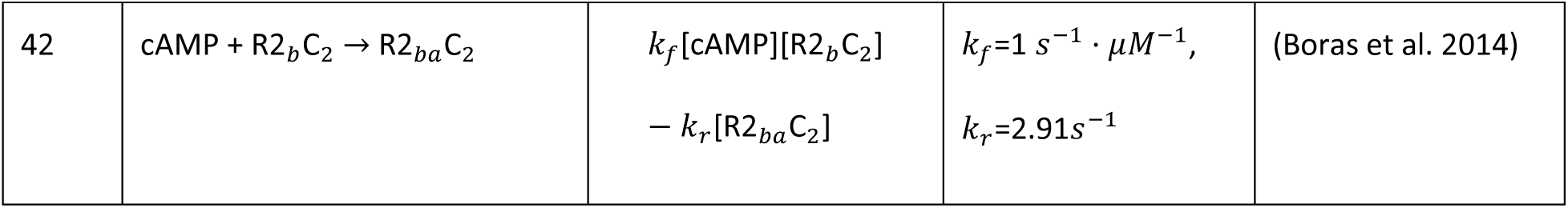
cAMP Reactions (cont.)

**Table S5:** Initial Conditions

| # | Species | Initial Value | Ref. |
| --- | --- | --- | --- |
| IC1 | AC8 | $1 \mu\text{M}$ | |
| IC2 | $\text{AC}_{ind}$ | $1 \mu\text{M}$ | |
| IC3 | $\text{Ca}^{2+}$ | $1 \mu\text{M}$ | |
| IC4 | CaM | $10 \mu\text{M}$ | |
| IC5 | cAMP | $0.1 \mu\text{M}$ | |
| IC6 | PDE1 | $1 \mu\text{M}$ | |
| IC7 | PDE4 | $0.4 \mu\text{M}$ | |
| IC8 | R2C2 | $0.4 \mu\text{M}$ | |
| IC9 | V | -60 mV |  |

### Minimal model to explore Ca^2+^-cAMP phase behavior

Based on the well-mixed model, we propose a minimal circuit to understand the phase behavior of Ca^2+^-cAMP. Consider the following system in **Figure 2c**, where Ca^2+^ is the stimulus which uses a pulse train to set influx times, AC is an activator, PDE1C is an inhibitor, and cAMP is the response element.

### Assumptions

The system is in a state such that the change in cAMP allows for further Ca^2+^ influx in a semi predictable manner, as represented by the pulse train. Therefore, this system is deemed to be stable if there exists a Ca^2+^ value that gives a stable solution for cAMP. All constants must be positive to remain physically relevant. We assume that there exists a constant independent source and sink within the system. The Ca^2+^ dependent and independent sources are localized homogeneously on the membrane, with both sinks located uniformly in the cytosol. For simplicity, we assume that the activation function for both *AC* and *PDE1C* are linear functions of Ca^2+^ of the form *aS* + *b*.

### Governing Equations

We define a stimulus (S, Ca^2+^) and a response element (R, cAMP). The well-mixed rate of change function for cAMP is then given by,

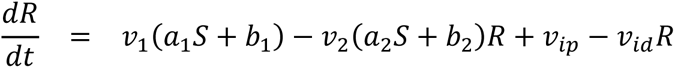

Here, *v*_1_ denotes the velocity of cAMP production by AC, *v*_2_ the rate of degradation by PDE, and *v_ip_* and *v_id_* the rates of independent production and degradation respectively.

### Analytical solutions for the minimal model

To analyze if the system lies in an in-or out-of-phase state we find the direction of the system change after initialization to *S*_0_ [*i.e.* the basal stimulus (initial concentration of Ca^2+^)]. First, we must solve for *R*_0_ (initial concentration of cAMP) by setting 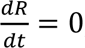, we find:

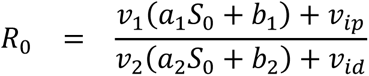

We start the system at equilibrium and prescribe a discontinuous pulse of *S* from *S*_0_ to *S*_ℎ_, akin to VGCC opening allowing Ca flux. Therefore, solving for the sign of R, we sobtain

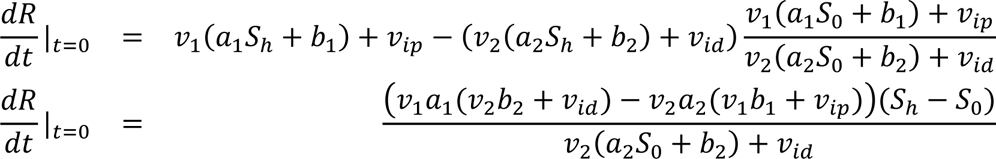

Since we consider the sign we can then characterize the solution by

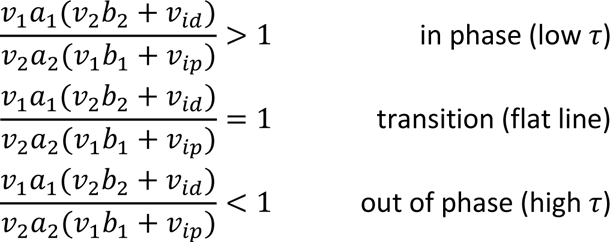

This model was then computationally run to confirm the results with arbitrary parameters (**Figure 2c**). The system was simulated with all parameters set to 1 except *v*_1_ and *v*_2_ at values of 4 and 2, respectively and vice-versa.

### Numerical implementation of the minimal model

We performed numerical simulations in MATLAB R2018b and checked against analytic solutions provided in the previous section. Minimal model solutions are found in **Figure 2c**.

### Simulations of the full spatial systems

For computational simplification, simulations were performed with a Gaussian profile on the top boundary, the size of the domain and Gaussian profile were informed by STORM images, **Figure 3**. The system is a hexagonal prism with outer radius of 0.4*μ*m and depth 0.6*μ*m with periodic boundary conditions in the x and y planes, see **Figure 4a**. The top plane is assumed to be the membrane and the bottom is a no flux condition. For the membrane plane, a Gaussian profile was normalized such that the average value is 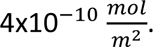 The Ca^2+^ sensitive AC initial conditions (Gaussian profile) is fixed for all simulation time by setting the diffusion constant to ≈0. Ca^2+^ sensitive AC initial conditions percent localization simulations move mass from the Gaussian profile into a uniform profile as a percent function (i.e a 25% Localization has 25% of the mass as a Gaussian profile and 75% as uniform). Examples of surface profiles at multiple % localizations can be seen in **Figure 4b**.

### Assumptions

- Membrane patterns are pre-existing and not effected by a single signaling event such that no diffusion occurs.
- Clustering events were approximated by a Gaussian profile in the center of the hexagonal prism.

### Model development

Although our system does show the ability to oscillate in and out of phase in a well-behaved manner, this will not produce our desired form if a homogenous boundary is present. Previous studies have looked at spatial gradients in the context of cAMP and PKA and found out that a localization must occur for the system to form one (Yang et al. 2016). Therefore, when moving to a 3D spatial map, we must consider how two solution regimes can be recovered.

Experimental data suggest that AKAP dimerizes and may form oligomers (Gold et al. 2011; Gao, Wang, and Malbon 2011), which is important for the function of these cells (Zhang et al. 2013). This could allow spatial instabilities like those seen in (Haselwandter et al. 2015) used to describe post synaptic domains. The final Gaussian profile on the top boundary had the size of the domain and Gaussian profile informed by STORM images of AC clustering (**Figure 3**).

Statistics of the images show that, on average, 90% of the AC in storm sits within 54 nm of the cluster center. The system was determined to be 0.35×0.35×0.6 *μ*m hexagonal prism with a Gaussian standard deviation of 25 nm.

Kinetic parameters were used the same as the well-mixed model, except for a few cases in which tuning through surface/volume relationships were needed. Post fitting was performed to further refine the relationship of PDE and AC through use of obtained FRET measurements (see section 4). Due to the large computational expense of the model PDE interactions were reduced to two steps (**Table S8**).

### Numerical Simulation

The well-mixed network was imported into COMSOL Multiphysics5.4 (Build:295), to solve the spatial model with in-homogeneous boundary conditions.

### Reaction-Diffusion Partial differential equations (PDEs)

Consider the same one-reaction system as the well-mixed system:

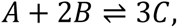

where the forward and backward rates are *k*_1_ and *k*_2_. But, now we spatially discretize the system, the partial differential equations describing the dynamics of species *A*, *B*, and *C* with unrestricted diffusion are then:

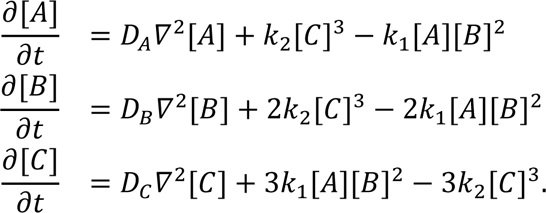

Where *∇*^2^ is the Laplacian and *D*_*A*_ is a diagonal matrix of diffusion coefficients for component A. Yet within the cell, all reactions do not occur in the free volume. In fact, most interactions occur on a membrane surface, which requires a different boundary condition.

### Reactions on the system boundary

Now let us assume the previous reaction occurs on the boundary and C is a membrane species on surface *Ω*

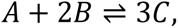

This would mean within the volume there is only free diffusion,

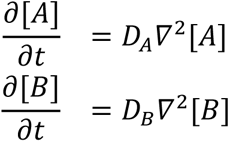

At the surface we would have a Neumann boundary condition, i.e a defined flux occurring into the boundary normal. We define the reaction as a flux occurring at a membrane surface (*Ω*)

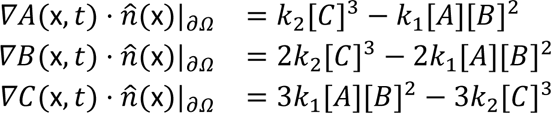

Here, *n̂* is the unit normal to *Ω*. We therefore have defined a flux between the volume and the membrane *Ω*. Finally, we consider *Ω* as a 2-dimensional surface existing at the system boundary with free diffusion of C;

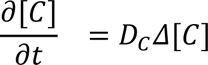

where *Δ*, is the surface Laplacian.

### PKA spatial response mirrors cAMP

PKA with full diffusion values 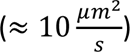 did not follow cAMP dynamics and only elicited one global response. We asked if this was due to AKAP patterning at the surface, which should mirror the AC profile at the surface. Adding this interaction into the model did not allow any sizable spatial gradient to develop. Recent work has suggested the PKA catalytic subunit in the presence of non-excess cAMP is effectively activated but its diffusion is restricted (F. Donelson Smith et al. 2017). By varying the diffusion constant, we found PKA activity could follow cAMP dynamics in the nanocluster and PM compartments in our computational model for restricted diffusivities (**Supplementary Fig. 5a**). We experimentally tested this prediction using the AKAP79-fused and PM-targeted PKA activity sensors and found that indeed PKA activity did follow the cAMP-Ca^2+^ phase relationship within the two compartments (**Supplementary Fig. 5b-d**). This suggests that anchored PKA holoenzyme action is much more restricted than originally anticipated.

### Comparisons with experimental data

Raw FRET data (**Figure 1**) was used for model refinement. The data was compared for oscillation time and phase, with expected cAMP concentration falling in the sensors sensitive range of ≈0.1-10*μ*M. Voltage gated channel sensitivities were not tuned, and only connection strengths between CaM to ACs and PDEs, which are largely less constrained in comparison, were varied. Values modified from the well-mixed model values can be found in **Tables S6-S9.**

### Model validation and predictions

The model was validated on predictions to concentration perturbations (AC, PDE, etc.) and disruption of patterning (AC binding disruption) and their changes to the phase of the signal. The system, once moved to the spatial model, was allowed free parameters along the sixsix-component axis for CaM connection and cAMP production strength of ACs and PDEs. This includes the flux differential between basal and activated ACs and PDEs.

### Reaction tables of modified well-mixed parameters for the spatial model

**Table S6:** Ca^2+^ Flux and reactions modified from Table S2.

| # | Reaction | Reaction Flux | Kinetic Parameters | Ref. |
| --- | --- | --- | --- | --- |
| S1 | $\rightarrow \text{Ca}$ | $j_{\text{Ca}_V}(1 + k_{\text{PKA}_V}[\text{PKA}])$<br>$+ j_{\text{Ca}_I}$ | $k_{\text{PKA}_V}=100\mu\text{M}^{-1}$ | FRET<br>constraint |
| S2 | $j_{\text{Ca}_V}$ | $f_i(-\alpha I_{\text{Ca}} - v_{\text{LPM}}[\text{Ca}])$ | $f_i=1\times 10^{-6}$, $\alpha=4.15\times 10^5$<br>$\text{mol} \cdot \text{m}^{-2} \cdot \text{A}^{-1} \cdot \text{s}^{-1}$,<br>$v_{\text{LPM}}=7.5\times 10^{-4} \text{m} \cdot \text{s}^{-1}$ | (Ni et al. 2011) |
| S3 | $j_{\text{Ca}_I}$ | $\frac{Ck_{\text{IP3R}}[\text{PKA}]A[\text{Ca}]}{1+A[\text{Ca}]+B[\text{Ca}]^2} ([\text{Ca}_{\text{stores}}] - [\text{Ca}]) - \frac{V_s[\text{Ca}]^2}{K_s^2+[\text{Ca}]^2}$ | $K_s=10\mu\text{M}$, $\text{Ca}_{\text{stores}}=1.56\mu\text{M}$,<br>$V_s=0.1\mu\text{M} \cdot \text{s}^{-1}$,<br>$k_{\text{IP3R}}=0.05\mu\text{M}^{-1}\text{s}^{-1}$,<br>$A=0.2869\mu\text{M}^{-1}$, $B=2.869\mu\text{M}^{-2}$,<br>$C=0.2133$ | (Ni et al. 2011) |
| S4 | $\text{AC} + \text{Ca}^2\text{CaM} \leftrightarrow \text{CaM} \cdot \text{AC}$ | $k_f[\text{AC}][\text{Ca}^2\text{CaM}]$<br>$- k_r[\text{CaM} \cdot \text{AC}]$ | $k_f=10.8 \text{s}^{-1} \cdot \mu\text{M}^{-1}$, $k_r=10 \text{s}^{-1}$ | FRET<br>constraint |
| S5 | $\text{CaM} \cdot \text{AC} + 2\text{Ca} \leftrightarrow \text{AC}^*$ | $K_{\text{cat}} \frac{[\text{Ca}][\text{CaM} \cdot \text{AC}]}{\text{Ca} + K_m}$<br>$- k_r[\text{AC}^*]$ | $K_{\text{cat}}=90 \text{s}^{-1}$, $K_m=1 \mu\text{M}$, $k_r=10 \text{s}^{-1}$ | FRET<br>constraint |
| S6 | $\text{PDE} + \text{Ca}^2\text{CaM} \leftrightarrow \text{CaM} \cdot \text{PDE}$ | $k_f[\text{PDE}][\text{Ca}^2\text{CaM}]$<br>$- k_r[\text{CaM} \cdot \text{PDE}]$ | $k_f=0.25 \text{s}^{-1} \cdot \mu\text{M}^{-1}$, $k_r=1 \text{s}^{-1}$ | FRET<br>constraint |
| S7 | $\text{CaM} \cdot \text{PDE} + 2\text{Ca} \leftrightarrow \text{PDE}^*$ | $K_{\text{cat}} \frac{[\text{Ca}][\text{CaM} \cdot \text{PDE}]}{\text{Ca} + K_m}$<br>$- k_r[\text{PDE}^*]$ | $K_{\text{cat}}=60 \text{s}^{-1}$, $K_m=1 \mu\text{M}$, $k_r=1 \text{s}^{-1}$ | FRET<br>constraint |
| S8 | $\text{PDE} + \text{Ca}^4\text{CaM} \leftrightarrow \text{PDE}^*$ | $k_f[\text{PDE}][\text{Ca}^4\text{CaM}]$<br>$- k_r[\text{PDE}^*]$ | $k_f=0.25 \text{s}^{-1} \cdot \mu\text{M}^{-1}$, $k_r=1 \text{s}^{-1}$ | FRET<br>constraint |

**Table S7:** cAMP reactions modified from Table S3.

| # | Reaction | Reaction Flux | Kinetic Parameters | Ref. |
| --- | --- | --- | --- | --- |
| S9 | → cAMP | $k_{base}([CaM \cdot AC] + [AC] + [AC_{ind}]) + k_{act}[AC^*]$ | $k_{base}=0.2 \text{ s}^{-1}$,<br>$k_{act}=23.55 \text{ s}^{-1}$,<br>$AC_{ind}=3 \times 10^{-8}$<br>$\text{mol} \cdot \text{m}^{-2}$ | FRET<br>constraint |
| S10 | cAMP→ | $k_{base}[cAMP] \frac{[CaM \cdot PDE] + [PDE]}{[cAMP] + K_m}$<br>$+ k_{act} \frac{[cAMP][PDE^*]}{[cAMP] + K_m}$ | $k_{base}=0.6 \text{ s}^{-1}$,<br>$K_m = 0.6 \text{ } \mu\text{M}$<br>$k_{act}=720 \text{ s}^{-1}$ | FRET<br>constraint |
| S11 | cAMP→ | $V_{ind} \frac{[cAMP]}{[cAMP] + K_m}$ | $V_{ind}=0.25 \text{ } \mu\text{M} \cdot \text{s}^{-1}$,<br>$K_m = 1.4 \text{ } \mu\text{M}$ | FRET<br>constraint |
| S12 | 2cAMP + R2C2 → R2C +<br>PKA | $k_f[cAMP]^2[R2C2] - k_r[R2C][PKA]$ | $k_f=20 \text{ min}^{-1} \cdot \mu\text{M}^{-2}$,<br>$k_r=12 \text{ min}^{-1} \cdot \mu\text{M}^{-1}$ | |
| S13 | 2cAMP + R2C → R2 +<br>PKA | $k_f[cAMP]^2[R2C] - k_r[R2][PKA]$ | $k_f=20 \text{ min}^{-1} \cdot \mu\text{M}^{-2}$,<br>$k_r=12 \text{ min}^{-1} \cdot \mu\text{M}^{-1}$ | |

**Table S8:** Added AKAP interactions for the spatial model.

| # | Reaction | Reaction Flux | Kinetic Parameters | Ref |
| --- | --- | --- | --- | --- |
| S14 | AKAP + R2 →<br>AKAP-R2 | $k_f[R2][AKAP-R2C2]$<br>$- k_r[AKAP-R2]$ | $k_f=1\text{ s}^{-1} \cdot \mu M^{-1}$<br>$k_r=0.1\text{ s}^{-1}$ | Est. |
| S15 | AKAP + R2C →<br>AKAP-R2C | $k_f[R2C][AKAP-R2C2]$<br>$- k_r[AKAP-R2C]$ | $k_f=1\text{ s}^{-1} \cdot \mu M^{-1}$<br>$k_r=0.1\text{ s}^{-1}$ | Est. |
| S16 | AKAP + R2C2 →<br>AKAP-R2C2 | $k_f[R2C2][AKAP-R2C2]$<br>$- k_r[AKAP-R2C2]$ | $k_f=1\text{ s}^{-1} \cdot \mu M^{-1}$<br>$k_r=0.1\text{ s}^{-1}$ | Est. |
| S17 | 2cAMP + AKAP-R2C2 → AKAP-R2C<br>+ PKA | $k_f[cAMP]^2[AKAP-R2C2]$<br>$- k_r[AKAP-R2C][PKA]$ | $k_f=20\text{ min}^{-1} \cdot$<br>$\mu M^{-2}, k_r=12$<br>$\text{min}^{-1} \cdot \mu M^{-1}$ | |
| S18 | 2cAMP + AKAP-R2C → AKAP-R2 +<br>PKA | $k_f[cAMP]^2[AKAP-R2C]$<br>$- k_r[AKAP-R2][PKA]$ | $k_f=20\text{ min}^{-1} \cdot$<br>$\mu M^{-2}, k_r=12$<br>$\text{min}^{-1} \cdot \mu M^{-1}$ | |

**Table S9:** ICs and diffusion.

| # | Species | Initial value | Diffusion | Ref. |
| --- | --- | --- | --- | --- |
| SIC1 | Ca<sup>2+</sup> | 0.001 $\mu M$ | 100 $\frac{\mu m^2}{s}$ | (Donahue and Abercrombie 1987), Est. from steady state |
| SIC2 | CaM | 2.9 $\mu M$ | 10 $\frac{\mu m^2}{s}$ | Est. from steady state<sup>1</sup> |
| SIC3 | Ca<sub>2</sub>CaM | 0.1 $\mu M$ | 10 $\frac{\mu m^2}{s}$ | Est. from steady state<sup>1</sup> |
| SIC4 | Ca<sub>3</sub>CaM | 4x10<sup>-3</sup> $\mu M$ | 10 $\frac{\mu m^2}{s}$ | Est. from steady state<sup>1</sup> |
| SIC5 | Ca<sub>4</sub>CaM | 1x10<sup>-2</sup> $\mu M$ | 10 $\frac{\mu m^2}{s}$ | Est. from steady state<sup>1</sup> |
| SIC6 | R<sub>2</sub> | 0.04 $\mu M$ | 10 $\frac{\mu m^2}{s}$ | Est. from steady state<sup>1</sup> |
| SIC7 | R<sub>2</sub>C<sub>2</sub> | 0.2 $\mu M$ | 10 $\frac{\mu m^2}{s}$ | Est. from steady state<sup>1</sup> |
| SIC8 | PKA | 0.05 $\mu M$ | 0.01 $\frac{\mu m^2}{s}$ | Est. from steady state, diffusion fitted |
| SIC9 | PDE1 | 0.9 $\mu M$ | 10 $\frac{\mu m^2}{s}$ | Est. from steady state<sup>1</sup> |
| SIC10 | PDE1<sub>act</sub> | 1x10<sup>-3</sup> $\mu M$ | 10 $\frac{\mu m^2}{s}$ | Est. from steady state<sup>1</sup> |
| SIC11 | CaMPDE1 | 1x10<sup>-3</sup> $\mu M$ | 10 $\frac{\mu m^2}{s}$ | Est. from steady state<sup>1</sup> |
| SIC12 | AC | 4x10<sup>-10</sup> $\frac{mol}{m^2}$ | 0 | Est. |
| SIC13 | CaMAC | 0 $\frac{mol}{m^2}$ | 0 | Est. |
| SIC14 | AC<sub>act</sub> | 0 $\frac{mol}{m^2}$ | 0 | Est. |
| SIC15 | AC<sub>ind</sub> | 4x10<sup>-10</sup> $\frac{mol}{m^2}$ | 1 $\frac{\mu m^2}{s}$ | Est.<sup>2</sup> |
| SIC16 | V | -60 mV | $1 \frac{\mu m^2}{s}$ | (Ni et al. 2011; Fridlyand, Tamarina, and Philipson 2003)<sup>2</sup> |
| SIC17 | cAMP | $4 \times 10^{-6} \mu M$ | $60 \frac{\mu m^2}{s}$ | (Agarwal, Clancy, and Harvey 2016), Est. from steady state |

<sup>1</sup>For all cytosolic species without well constrained diffusions we defined them as $10 \frac{\mu m^2}{s}$
<sup>2</sup>For all membrane species without well constrained diffusions we defined them as $1 \frac{\mu m^2}{s}$

---

**Extended Figure 1.**
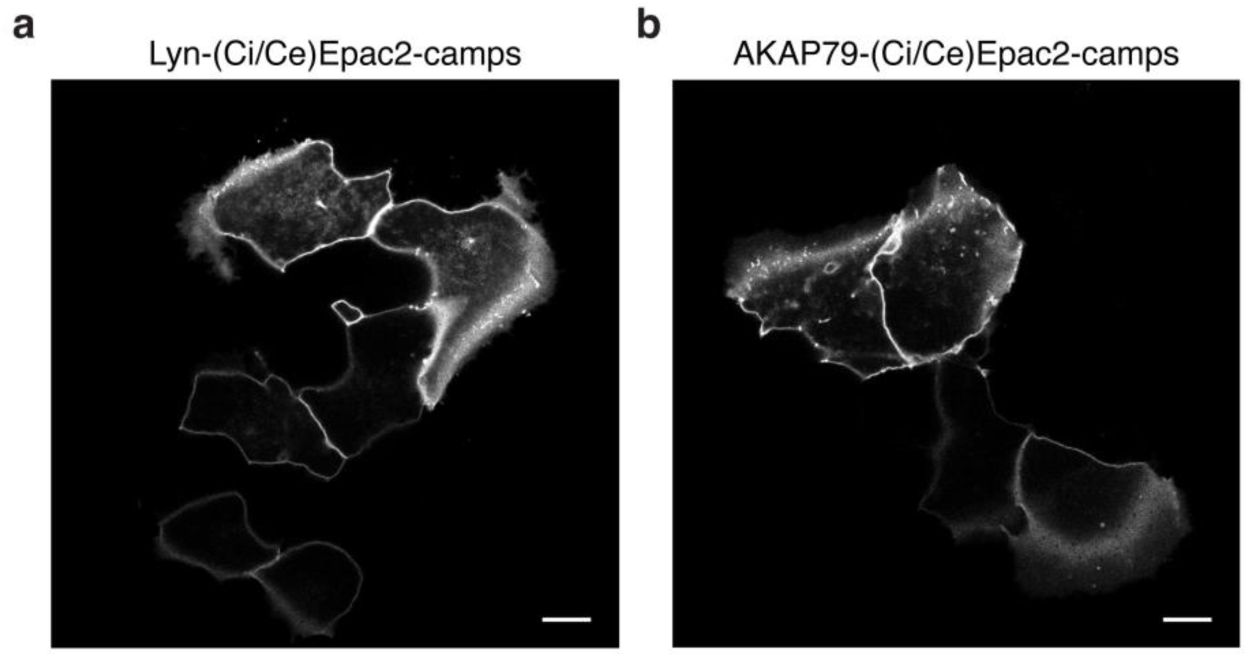
The AKAP79/150 and PM targeted sensors are localized at the plasma membrane. (a) Confocal image of lyn(Ci/Ce)Epac2camps showing efficient localization of the probe at the PM in MIN6 cells (YFP channel, scale 5μm). (b) Confocal image of AKAP79-(Ci/Ce)Epac2camps also depicting localization of the scaffold-fused biosensor at the PM in MIN6 cells (YFP channel, scale 5μm).

**Extended Figure 2.**
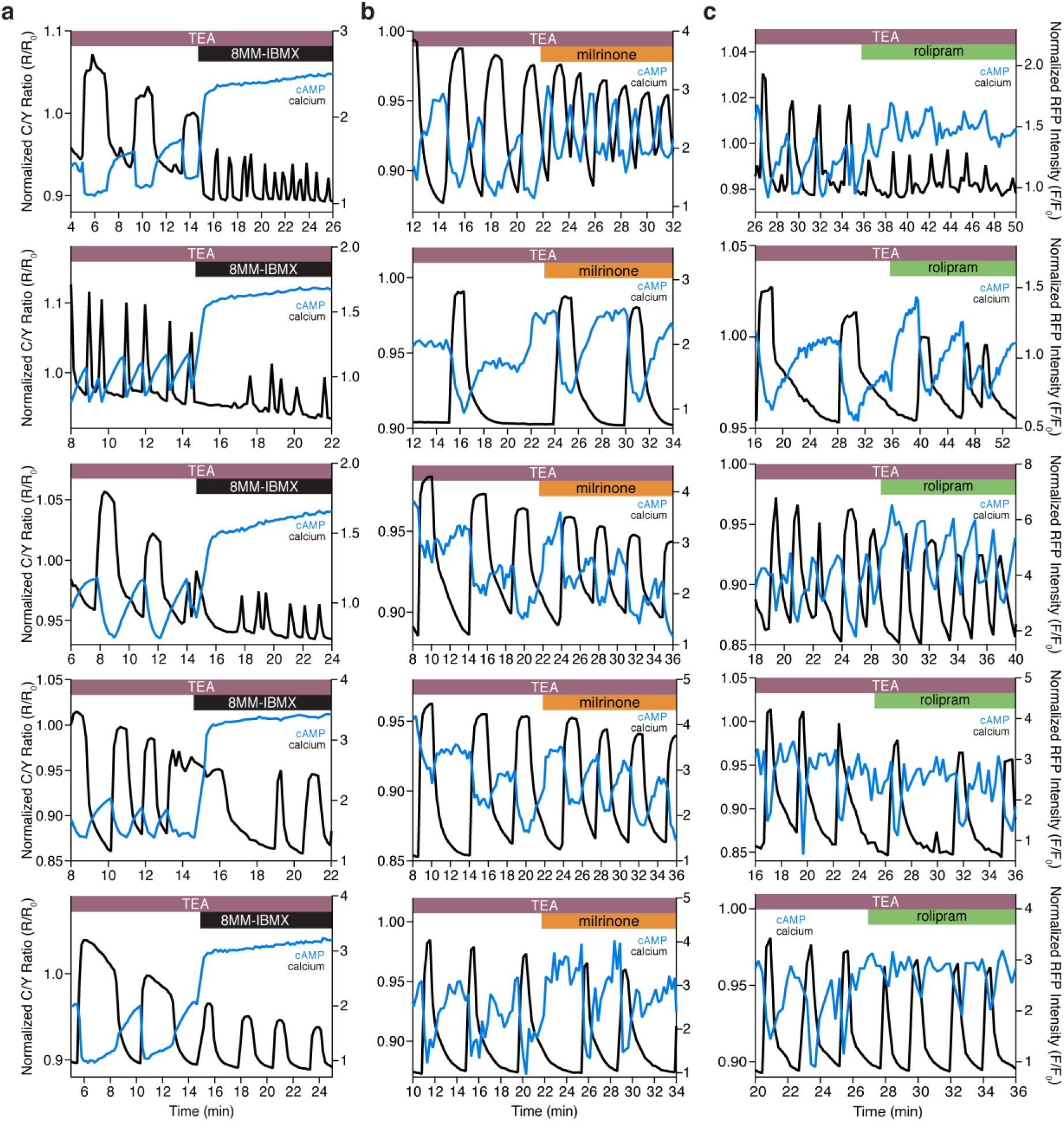
The Ca^2+^-dependent cAMP response is dependent on PDE1, but not PDE3 or PDE4. (a) Representative single cell traces depicting the decoupling of oscillating cAMP at the PM (measured by lyn-(Ci/Ce)Epac2camps, blue trace) from oscillating Ca^2+^ (black trace) upon inhibition of PDE1C with 8MM-IBMX. Representative single cell traces showing cAMP still oscillates at the PM (blue trace) together with Ca^2+^ (black trace) upon inhibition of PDE3 (milrinone) (b) and PDE4 (rolipram) (c), respectively.

**Extended Figure 3.**
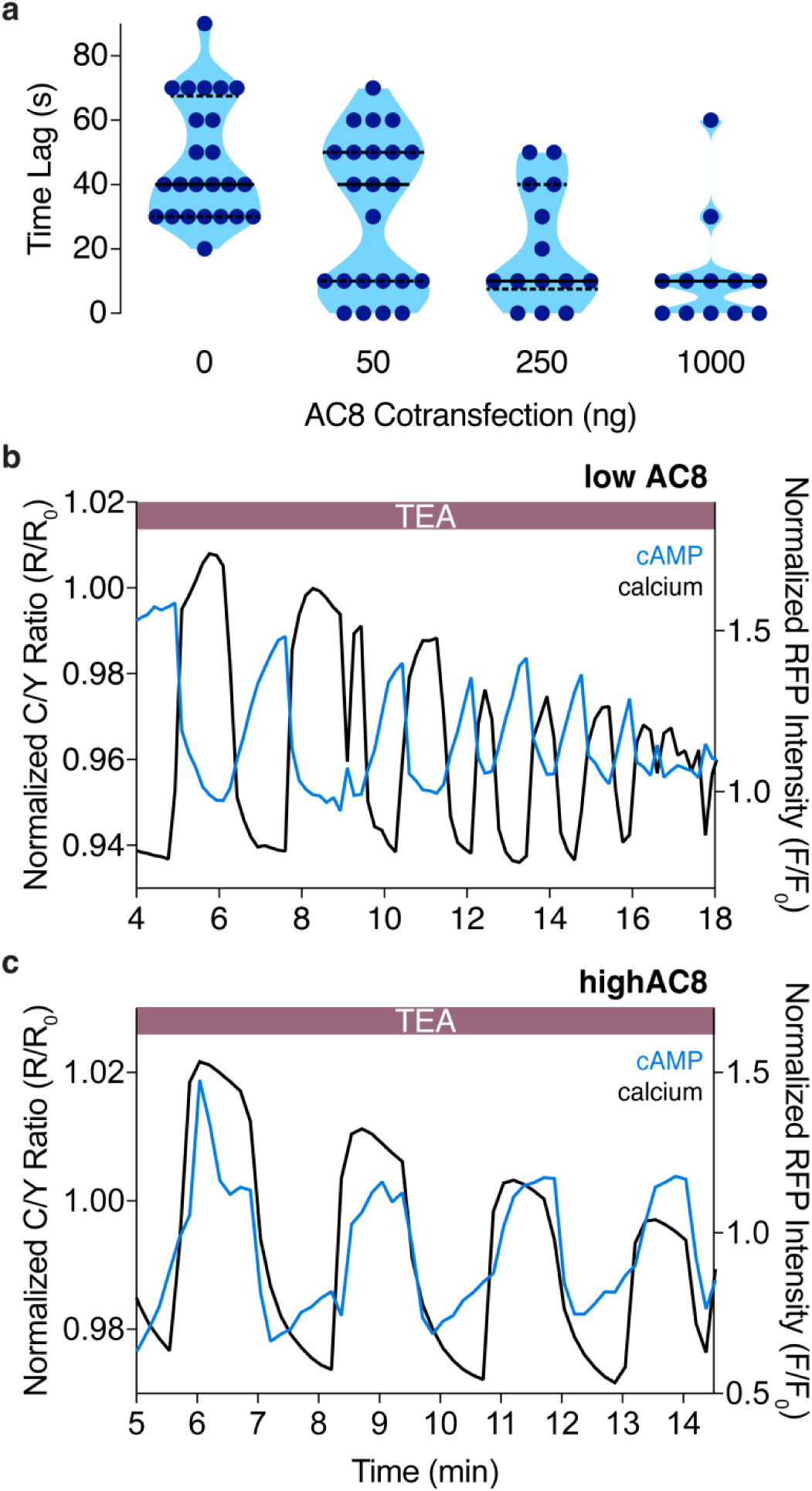
The cAMP-Ca^2+^ phase relationship at the PM can be tuned by expression of the Ca^2+^-dependent AC8. (a) Plot depicting a dose-dependence between the time lag of cAMP at the PM and the amount of AC8 co-transfected (n = 56). (b) Representative single cell trace of an oscillating β cell control with no AC8 co-expression, illustrating an out-of-phase cAMP-Ca^2+^ phase relationship. Blue trace is cAMP at PM and black trace is Ca^2+^. (c) Representative single cell trace of an oscillating β cell with 1μg of an AC8 expression vector co-transfected, illustrating an in-phase cAMP-Ca^2+^ phase relationship. Blue trace is cAMP at PM and black trace is Ca^2+^.

**Extended Figure 4.**
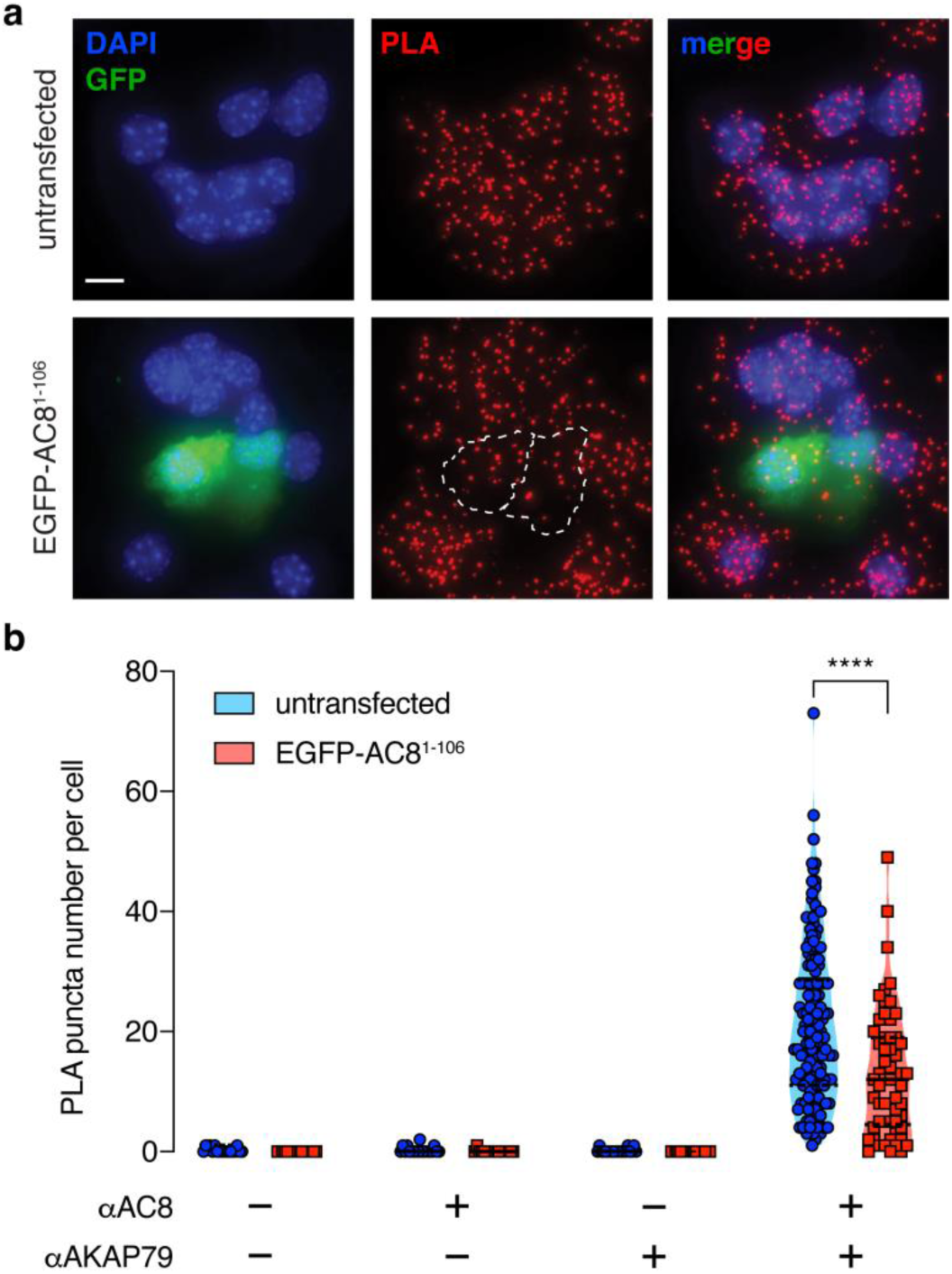
Expression of the N-terminus of AC8 is sufficient to perturb the interaction between endogenous AKAP150 and AC8 in MIN6. (a) Widefield maximum intensity projections of a Proximity Ligation Assay to depict the extent of interactions between AC8 and AKAP150 in MIN6 β cells (scale 10 μm). Expression of EGFP-tagged AC8^1-106^ results in less PLA puncta per cell. (b) Number of PLA puncta per AC8^1-106^-expressing cell is significantly decreased compared to non-transfected control (n = 142 and 57 cells, respectively) (p<0.05).

**Extended Figure 5.**
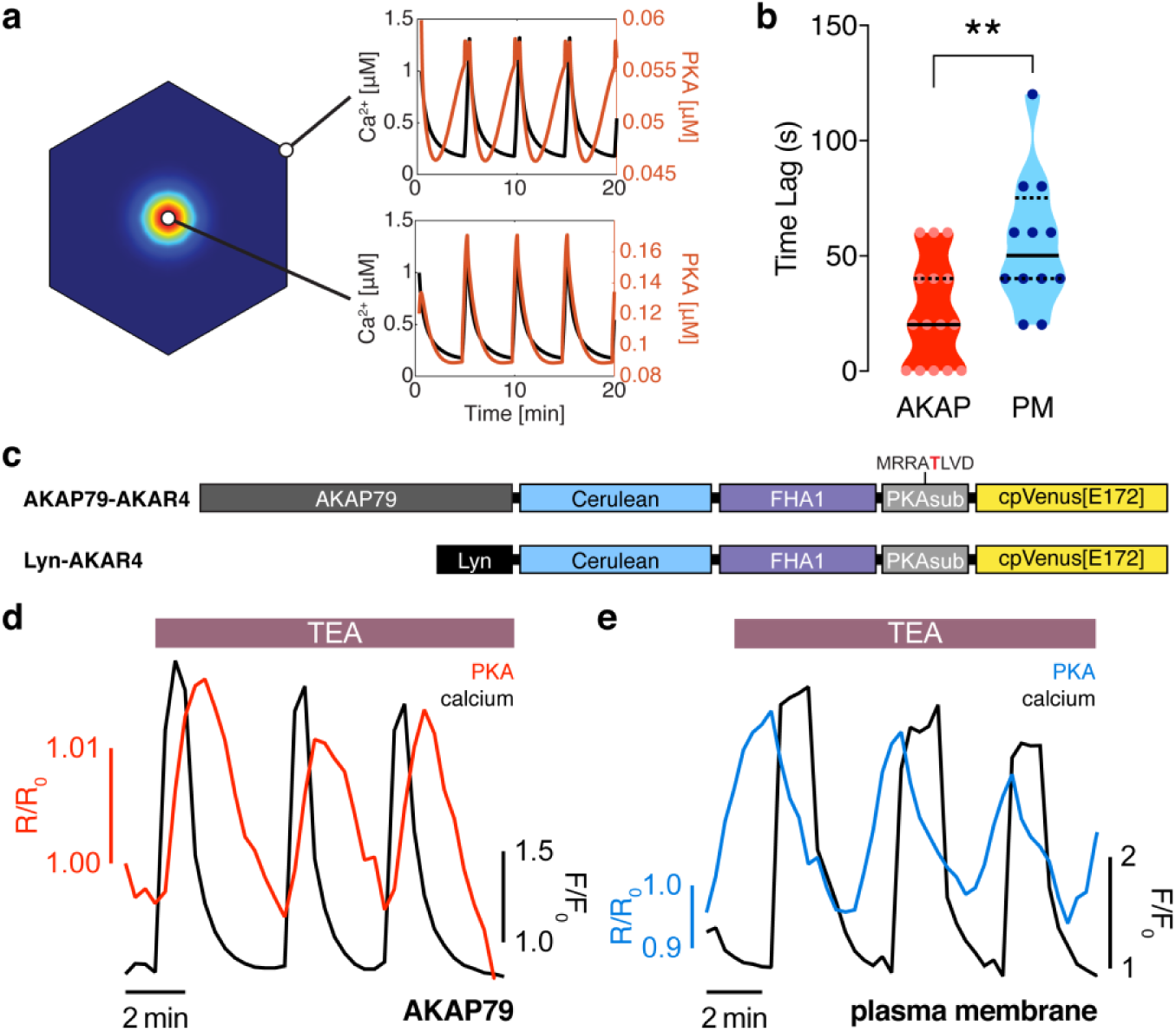
Oscillating PKA activity phase is also spatially compartmentalized at the PM. (a) PKA activity is predicted to oscillate near the center of the AKAP79/150:AC8 cluster with a short time delay and a longer delay outside of the cluster, relative to Ca^2+^. (b) Experimentally-measured time lag between PKA activity oscillations and Ca^2+^ for AKAP79 (orange) and the general PM (teal) compartments (n=15 and 12, respectively). (c) Representative single cell trace showing in-phase PKA activity within the AKAP79 compartment. Orange trace is PKA activity at AKAP79 (CY-FRET channel divided by cyan donor channel) and black trace is Ca^2+^ (RFP). (d) Representative single cell trace showing out-of-phase PKA activity within the general PM compartment. Teal trace is PKA activity at PM (CY-FRET channel divided by cyan donor channel) and black trace is Ca^2+^ (RFP).

**Accessory Figure 1.**
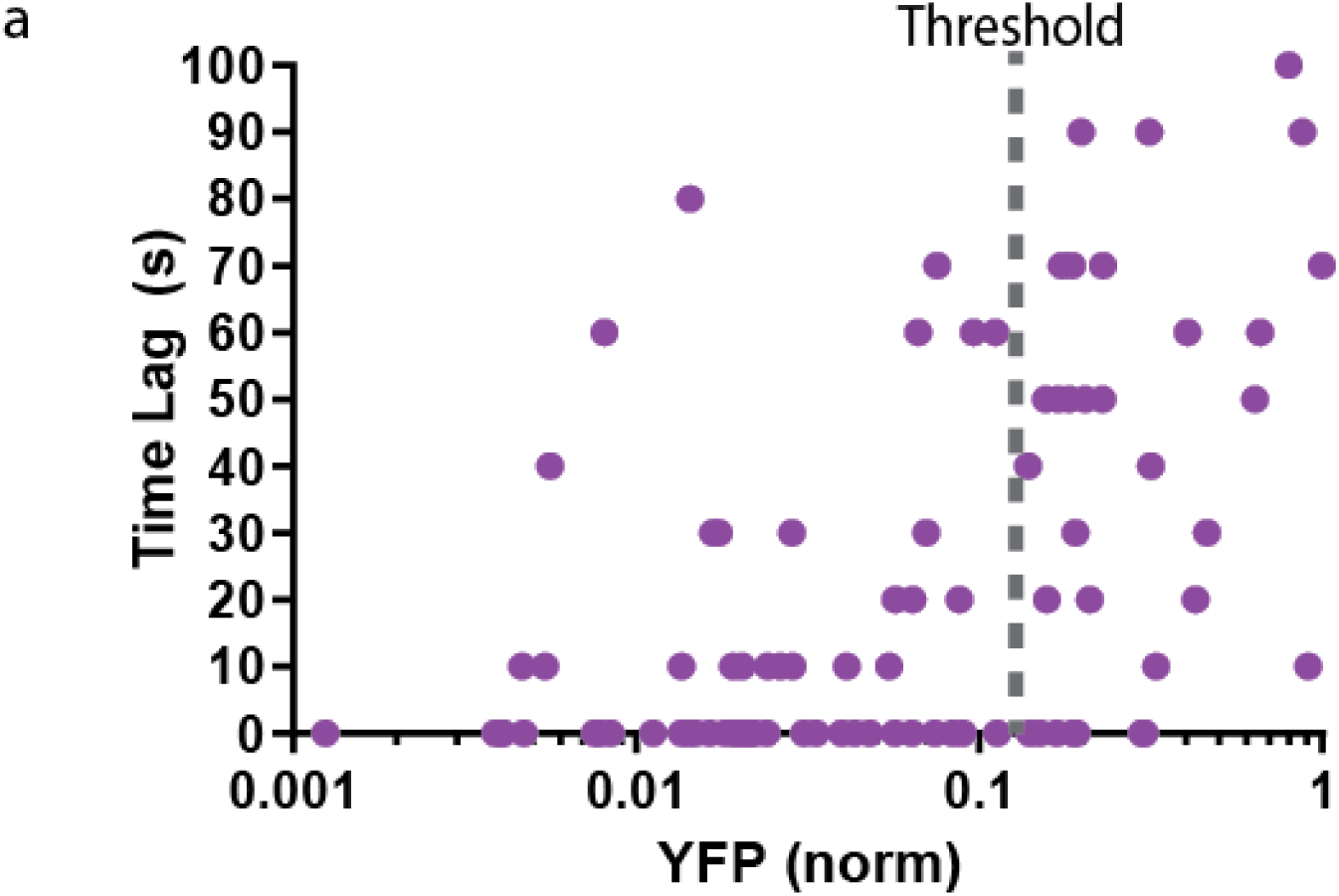
Phase of cAMP correlates with expression level of the AKAP79-(Ci/Ce)Epac2-camps. (a) Scatter plot of the time lag (sec) and the YFP donor channel intensity (normalized to non-saturating maximum) for each cell expressing AKAP79-(Ci/Ce)Epac2camps. Cells with higher expression of the probe correlated with a longer time lag, therefore a YFP intensity threshold was designated for analysis purposes.

**Accessory Figure 2.**
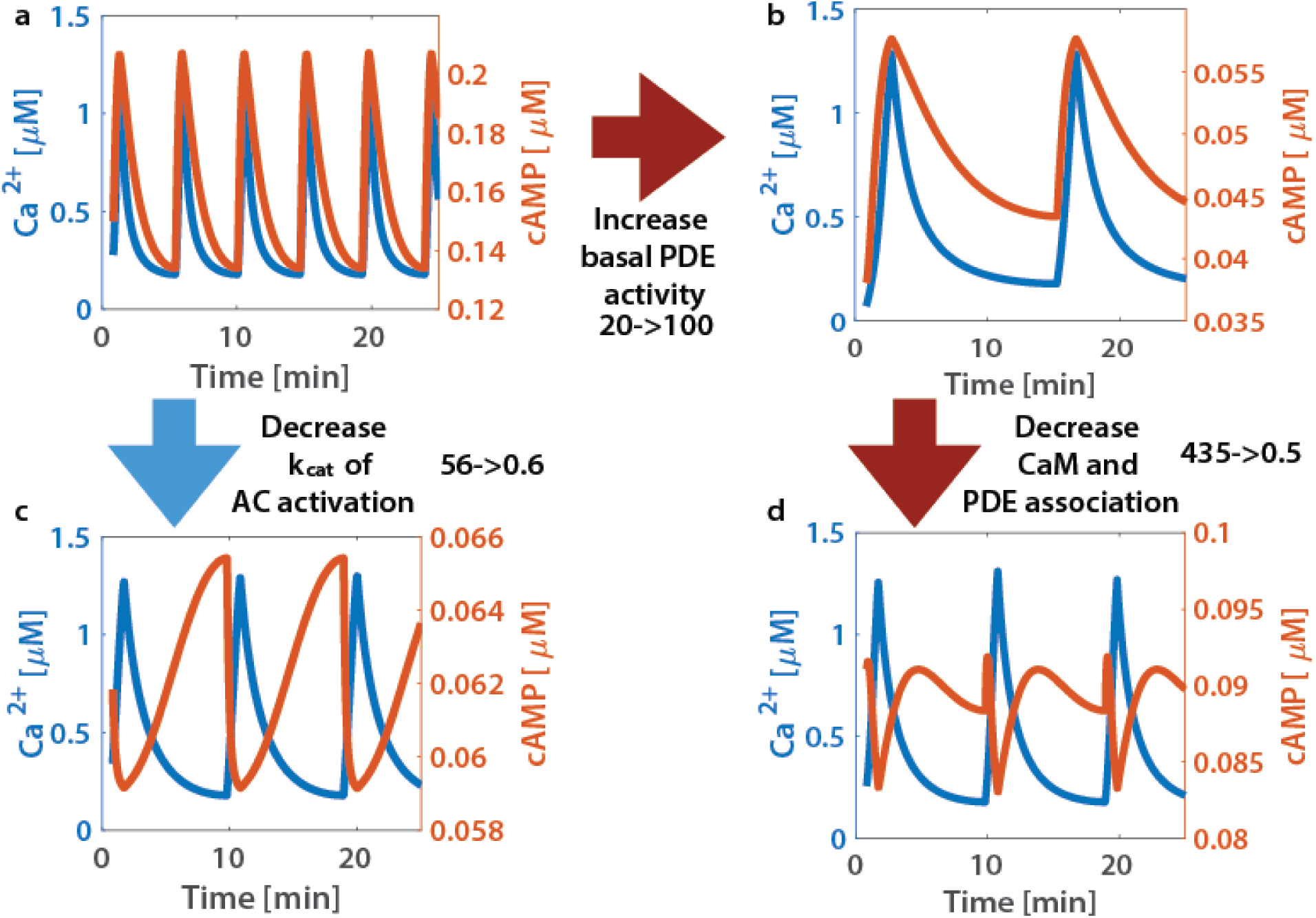
**Phase is driven by activity variability within the Ca**^2+^ **oscillatory regime.** System phase can be switched by tuning the association of CaM to sources (ACs) and sinks (PDEs). (**a**) At base system conditions, the system acts in an in-phase manner. (**c**) Decreasing the rate of Ca^2+^ association to the AC-CaM complex causes the phase to switch to out-of-phase. (**b**) increasing basal PDE activity does not allow a phase switch, only after decreasing PDE and CaM association rates will the system allow a phase switch (**c**). A phase switch is controlled by the variability in the activity of source or sink. If the sink dominates, then the system is out of phase. If the source dominates, the system is in phase.

## References

1. Agarwal, SR. et al. (2016). “Mechanisms restricting diffusion of intracellular camp.” Sci. Rep. 6: 19577.

2. Akerboom, J. et al. (2013). “Genetically encoded calcium indicators for multi-color neural activity imaging and combination with optogenetics.” Front. Mol. Neurosci. 6:2.

3. Ang, K. et al. (2002). “Reciprocal regulation of calcium dependent and calcium independent cyclic AMP hydrolysis by protein phosphorylation.” J. of Neurochemistry. 81(3):422:433.

4. Barg, S. et al. (2001). “Fast Exocytosis with Few Ca^2+^ Channels in Insulin-secreting Mouse Pancreatic Beta Cells.” Biophysical Journal. 81(6): 3308–3323.

5. Beene, D. et al. (2007). “A-kinase chaoring proteins take shape.” Curr. Opin. Cell. Biol. 19(2): 192–198.

6. Bender, AT. et al. (2006). “Cyclic nucleotide phosphodiesterases: molecular regulation to clinical use.” Pharmacol Rev. 58(3): 488–520.

7. Berridge, M. et al. (1998). “Calcium – a life and death signal.” Nature 395 645–648.

8. Bertram, R. et al. (2011). “Electrical Bursting, Calcium Oscillations, and Synchronization of Pancreatic Islets.” Adv Exp Med Biol. 654: 261–279.

9. Bohrer et al. (2019). “A Pairwise Distance Distribution Correction (DDC) algorithm for blinking-free and super-resolution microscopy.” bioRxiv.

10. Calebiro, D. et al. (2014). “cAMP signaling microdomains and their observation by optical methods.” Front. Cell. Neurosci.

11. Clapham, D. (2007). “Calcium Signaling.” Cell 131(6): 1047–1058.

12. Conti, M. et al. (2014). “Cyclic AMP compartments and signaling specificity: role of cyclic nucleotide phophodiesterases.” J. Gen. Physiol. 143: 29–38.

13. Cooper, DM. et al. (1995). “Adenylyl cyclases and the interaction between calcium and cAMP signalling.” Nature. 374(6521): 421–4.

14. De Pitta, M. et al. (2008). “Coexistence of amplitutde and frequency modulations in intracellular calcium dynamics.” Physical Review E 77, 030903(R).

15. Defer, N. et al. (2000). “Tissue specificity and physiological relevance of various isoforms of adenylyl cyclase.” American Journal of Physiology – Renal Physiology. 279(3): F400–F416.

16. Delint-Ramirez, I. et al. (2011). “Palmitoylation targets AKAP79 protein to lipid rafts and promotes its regulation of calcium-sensitive adenylyl cyclase type 8.” J. Biol. Chem. 286(38): 32962–75.

17. Dell’Acqua, ML. et al. (2006). “Regulation of neuronal PKA signaling through AKAP targeting dynamics.” Eur J Cell Biol. 85(7):627–33.

18. Depry, C. et al. (2011). “Visualization of PKA activity in plasma membrane microdomains.” Mol Biosyst. 7(1): 52–8.

19. Dou, H. et al. (2015). “Calcium influx activates adenylyl cyclase 8 for sustained insulin secretion in rat pancreatic beta cells.” Diabetologia. 58(2): 324–333.

20. Draznin, B. (1988) “Intracellular calcium, insulin secretion, and action.” Am J Med. 85(5A): 44–58.

21. Dupont, G. et al. (2011). “Calcium Oscillations.” Cold Spring Harb Perspect Biol 3(3).

22. Dyachok, O. et al. (2006). “Oscillations of cyclic AMP in hormone-stimulated insulin-secreting beta-cells.” Nature 439(7074): 349–52.

23. Everett, E. et al. (2013). “An Improved Targeted cAMP Sensor to Study the Regulation of Adenylyl Cyclase 8 by Ca^2+^ Entry through Voltage-Gated Channels.” PLoS One. 8(9): e75942.

24. Fridlyand, LE. et al. (2007). “Regulation of cAMP dynamics by Ca^2+^ and G protein-coupled receptors in the pancreatic beta cell: a computational approach.” Am J Physiol Cell Physiol. 293(6):C1924–33.

25. Fridlyand, LE. et al. (2010). “Bursting and calcium oscillations in pancreatic beta-cells: specific pacemakers for specific mechanisms.” Am J Physiol Endocrinol Metab. 299(4): E517–32.

26. Gao, J. et al. (2016). “Differential role of SNAP-25 phosphorylation by protein kinases A and C in the regulation of SNARE complex formation and exocytosis in PC12 cells.” Cell Signaling. 28(5): 425–437.

27. Gilon, P. et al. (2002). “Control Mechanisms of the Oscillations of Insulin Secretion In Vitro and In Vivo.” Diabetes. 51:S144–S151.

28. Gold, M. et al. (2011). “Architecture and dynamics of an A-kinase anchoring protein 79 (AKAP79) signaling complex.” PNAS. 108(16): 6426–6431.

29. Goraya, TA. et al. (2008). “Kinetic properties of Ca^2+^/calmodulin-dependent phosphodiesterase isoforms dictate intracellular cAMP dynamics in response to elecation of cytosolic Ca^2+^.” Cell Signaling. 20(2): 359–74.

30. Greenwald, E. et al. (2011). “Bigger, better, faster: principles and models of AKAP signaling.” J. Cardiobasc. Pharmacol. 58(5): 462–469.

31. Gromada, J. et al. (2004). “Glucagon-Like Peptide-1: Regulation of Insulin Secretion and Therapeutic Potential.” Basic and Clinical Pharmacology and Toxicology. 95(6).

32. Gucek, A. et al. (2018). “Fusion pore regulation by Epac2/cAMP controls cargo release during insulin exocytosis.” bioRxiv.

33. Han, P. et al. (1999). “The calcium/calmodulin-dependent phosphodiesterase PDE1C down-regulates glucose-induced insulin secretion.” J. Biol. Chem. 273(32): 22337–44.

34. Hanoune, J. et al. (2001). “Regulation and Role of Adenylyl Cyclase Isoforms.” Annual Review of Pharmacology and Toxicology. 41: 145–174.

35. Hinke, S. et al. (2012). “Anchored phosphatases modulate glucose homeostasis.” EMBO Journal. 31(20): 3991–4004.

36. Landa, L. et al. (2005). “Interplay of Ca^2+^ and cAMP Signaling in the Inulin-secreting MIN6 Beta-Cell Line.” JBC. 280, 31294–31302.

37. Levchenko, A. et al. (2000). “Scaffold proteins may biphasically affect the levels of mitogen-activated protein kinase signaling and reduce its threshold properties.” PNAS. 97(11): 5818-5823.

38. Lohse, C. et al. (2017). “Experimental and mathematical analysis of cAMP nanodomains.” PLoS One.

39. Masada, N. et al. (2008). “Distinct Mechanisms of Regulation by Ca^2+^/Calmodulin of Type 1 and 8 Adenylyl Cyclases Support Their Different Physiological Roles.” JBC. 284:4451–4463.

40. Masada, N. et al. (2012). “Distinct Mechanisms of Calmodulin Binding and Regulation of Adenylyl Cyclases 1 and 8.” Biochemistry. 51, 40, 7917–7929.

41. Mo, GC. et al. (2017). “Genetically encoded biosensors for visualizing live-cell biochemical activity at super-resolution.” Nature Methods. 14(4): 427–434.

42. Murphy, J. et al. (2014). “AKAP-Anchored PKA Maintains Neuronal L-type Calcium Channel Activity and NFAT Transcriptional Signaling.” Cell Rep. 7(5): 1577–1588.

43. Musheshe, N. et al. (2018). “cAMP: From Long-Range Second Messenger to Nanodomain Signalling.” Trends in Pharmacological Sciences. 39(2): 209–222.

44. Musheshe, N. et al. (2018). “Targeting FRET-Based Reporters for cAMP and PKA Activity Using AKAP79.” Sensors (Basel). 18(7): pii: E2164.

45. Nesher, R. et al. (2002). “Beta-cell protein kinases and the dynamics of the insulin response to glucose.” Diabetes. 51 Suppl. 1:S68–73.

46. Ni, Q. et al. (2011) “Signaling Diversity of PKA Achieved Bia a Ca^2+^-cAMP-PKA Oscillatory Circuit.” Nat Chem Biol. 7(1): 34–40.

47. Oliveria, S. et al. (2003). “Imaging kinase-AKAP79-phoshpatase scaffold complexes at the plasma membrane in living cells using FRET microscopy.” J Cell Biol. 160(1): 101–112.

48. Parekh, A. (2010). “Decoding cytosolic Ca^2+^ oscillations.” Trends in Biochemical Sciences. 36(2): P78–87.

49. Peercy, B. et al. (2015). “Modeling of Glucose-Induced cAMP Oscillations in Pancreatic Beta Cells: camp Rocks when Metabolism Rolls.” Biophysical Journal. 109(2): 439–449.

50. Peterson, OH. (2002). “Calcium signal compartmentalization.” Biol. Res. 35(2): 177–82.

51. Purkey, AM. et al. (2018). “AKAP150 Palmitoylation Regulates Synaptic Incorporation of Ca^2+^-Permeable AMPA Receptors to Control LTP.” Cell Rep. 25(4): 974–987.

52. Purvis, J. et al. (2014). “Encoding and Decoding Cellular Information through Signaling Dynamics.” Cell. 152(5): 945–956.

53. Raoux, M. et al. (2015). “Multilevel control of glucose homeostasis by adenylyl cyclase 8.” Diabetologia. 58(4): 749–757.

54. Renstrom, E. et al. (2004). “Protein kinase A-dependent and –independent stimulation of exocytosis by cAMP in mouse pancreatic beta-cells.” Journal of Physiology. 502(1).

55. Rorsman, P. et al. (2018). “Pancreatic β-Cell Electrical Activity and Insulin Secretion: Of Mice and Men.” Physiol. Rev. 98: 117–214.

56. Sassone-Corsi, P. (2012). “The Cyclic AMP Pathway.” Cold Spring Harb Perspect Biol 4(12).

57. Saucerman, J. et al. (2013). “Mechanisms of cyclic AMP compartmentation revealed by computational models.” J. Gen. Physiol. 143(1): 39–48.

58. Schmitz, O. et al. (2002). “High-frequency insulin pulsatility and type 2 diabetes: from physiology and pathophysiology to clinical pharmacology.” Diabetes Matabolism. 28(6 Suppl): 4S14–20.

59. Smith, FD. et al. (2002). “Signaling complexes: junction on the intracellular information super highway.” Curr. Biol. 12(1):R32–40.

60. Stern, MD. (1992). “Buffering of calcium in the vicinity of a channel pore.” Cell Calcium. 13(3): 183–92.

61. Tajada, S. et al. (2017). “Distance constraints on activation of TRPV4 channels by AKAP150-bound PKCalpha in arterial myocytes.” JGP. 149(6): 639.

62. Tengholm, A. (2012). “Cyclic AMP dynamics in the pancreatic β-cell.” Ups. J. Med. Sci. 117(4): 355–69.

63. Thompson, JL. et al. (2015). “Anchoring protein AKAP79-mediated PKA phosphorylation of STIM1 determines selective activation of the ARC channel, a store-independent Orai channel.” J Physiol. 593(3): 559–72.

64. Torres-Quesada, O. et al. (2017). “The many faces of compartmentalized PKA signalosomes.” Cellular Signalling. 37: 1–11.

65. Wachten, S. et al. (2010). “Distinct pools of cAMP centre on different isoforms of adenylyl cyclase in pituitary-derived GH_3_B_6_ cells.” J Cell Sci. 123: 95–106.

66. White, MA. et al. (2005). “Signaling networks in living cells.” Annu. Rev. Pharmacol. Toxicol. 45: 587–603.

67. Willoughby, D. et al. (2006). “Ca^2+^ stimulation of adenylyl cyclase generates dynamic oscillations in cyclic AMP.” Journal of Cell Science. 119:828–836.

68. Willoughby, D. et al. (2010). “AKAP79/150 Interacts with AC8 and Regulates Ca^2+^-dependent cAMP Synthesis in Pancreatic and Neuronal Systems.” J Biol Chem. 285(26): 20328–20342.

69. Willoughby, D. et al. (2012). “A key phosphorylation site in AC8 mediates regulation of Ca^2+^-dependent cAMP dynamics by an AC8-AKAP79-PKA signaling complex.” J Cell Sci. 125(23): 5850–5859.

70. Zhang, J. et al. (2016). “Clustering and Functional Coupling of Diverse Ion Channels and Signaling Proteins Revealed by Super-resolution STORM Microscopy in Neurons.” Neuron. 92(2): 461–478.

71. Zhang, X. et al. (2019). “A calcium/cAMP signaling loop at the ORAI1 mouth drives channel inactivation to shape NFAT induction.” Nature Communications. 10: 1971.

## Supplementa References

72. Agarwal, Shailesh R., Colleen E. Clancy, and Robert D. Harvey. 2016. “Mechanisms Restricting Diffusion of Intracellular cAMP.” Scientific Reports 6 (January): 19577. https://doi.org/10.1038/srep19577.

73. Ang, Kok-Long, and Ferenc A. Antoni. 2002. “Reciprocal Regulation of Calcium Dependent and Calcium Independent Cyclic AMP Hydrolysis by Protein Phosphorylation.” Journal of Neurochemistry 81 (3): 422–33. https://doi.org/10.1046/j.1471-4159.2002.00903.x.

74. Berry, Hugues. 2002. “Monte Carlo Simulations of Enzyme Reactions in Two Dimensions: Fractal Kinetics and Spatial Segregation.” Biophysical Journal 83 (4): 1891–1901. https://doi.org/10.1016/S0006-3495(02)73953-2.

75. Boras, Britton W., Alexandr Kornev, Susan S. Taylor, and Andrew D. McCulloch. 2014. “Using Markov State Models to Develop a Mechanistic Understanding of Protein Kinase A Regulatory Subunit RI*α* Activation in Response to cAMP Binding.” Journal of Biological Chemistry 289 (43): 30040–51. https://doi.org/10.1074/jbc.M114.568907.

76. Donahue, B. S., and R. F. Abercrombie. 1987. “Free Diffusion Coefficient of Ionic Calcium in Cytoplasm.” Cell Calcium 8 (6): 437–48. https://doi.org/10.1016/0143-4160(87)90027-3.

77. Fridlyand, Leonid E., and Louis H. Philipson. 2011. “Coupling of Metabolic, Second Messenger Pathways and Insulin Granule Dynamics in Pancreatic Beta-Cells: A Computational Analysis.” *Progress in Biophysics and Molecular Biology*, Beta-cell simulation and Clinical studies on insulin secretion, 107 (2): 293–303. https://doi.org/10.1016/j.pbiomolbio.2011.09.001.

78. Fridlyand, Leonid E., and Louis H. Philipson. 2016. “Pancreatic Beta Cell G-Protein Coupled Receptors and Second Messenger Interactions: A Systems Biology Computational Analysis.” PLOS ONE 11 (5): e0152869. https://doi.org/10.1371/journal.pone.0152869.

79. Fridlyand, Leonid E., Natalia Tamarina, and Louis H. Philipson. 2003. “Modeling of Ca2+ Flux in Pancreatic Beta-Cells: Role of the Plasma Membrane and Intracellular Stores.” American Journal of Physiology. Endocrinology and Metabolism 285 (1): E138–154. https://doi.org/10.1152/ajpendo.00194.2002.

80. Gao, Shujuan, Hsien-yu Wang, and Craig C. Malbon. 2011. “AKAP12 and AKAP5 Form Higher-Order Hetero-Oligomers.” Journal of Molecular Signaling 6 (1): 8. https://doi.org/10.1186/1750-2187-6-8.

81. Gold, Matthew G., Florian Stengel, Patrick J. Nygren, Chad R. Weisbrod, James E. Bruce, Carol V. Robinson, David Barford, and John D. Scott. 2011. “Architecture and Dynamics of an A-Kinase Anchoring Protein 79 (AKAP79) Signaling Complex.” Proceedings of the National Academy of Sciences 108 (16): 6426–31. https://doi.org/10.1073/pnas.1014400108.

82. Guldberg, C. M., and P. Waage. 1879. “Ueber Die Chemische Affinität. 1. Einleitung.” Journal Für Praktische Chemie 19 (1): 69–114. https://doi.org/10.1002/prac.18790190111.

83. Han, Ping, John Werber, Manju Surana, Norman Fleischer, and Tamar Michaeli. 1999. “The Calcium/Calmodulin-Dependent Phosphodiesterase PDE1C down-Regulates Glucose-Induced Insulin Secretion.” Journal of Biological Chemistry 274 (32): 22337–44. https://doi.org/10.1074/jbc.274.32.22337.

84. Haselwandter, Christoph A., Mehran Kardar, Antoine Triller, and Rava Azeredo da Silveira. 2015. “Self-Assembly and Plasticity of Synaptic Domains Through a Reaction-Diffusion Mechanism.” Physical Review E 92 (3): 032705. https://doi.org/10.1103/PhysRevE.92.032705.

85. Lai, Massimo, Denis Brun, Stuart J. Edelstein, and Nicolas Le Novère. 2015. “Modulation of Calmodulin Lobes by Different Targets: An Allosteric Model with Hemiconcerted Conformational Transitions.” PLOS Computational Biology 11 (1): e1004063. https://doi.org/10.1371/journal.pcbi.1004063.

86. Masada, Nanako, Antonio Ciruela, David A. MacDougall, and Dermot M. F. Cooper. 2009. “Distinct Mechanisms of Regulation by Ca2+/Calmodulin of Type 1 and 8 Adenylyl Cyclases Support Their Different Physiological Roles.” Journal of Biological Chemistry 284 (7): 4451–63. https://doi.org/10.1074/jbc.M807359200.

87. Masada, Nanako, Sabine Schaks, Sophie E. Jackson, Andrea Sinz, and Dermot M. F. Cooper. 2012. “Distinct Mechanisms of Calmodulin Binding and Regulation of Adenylyl Cyclases 1 and 8.” Biochemistry 51 (40): 7917–29. https://doi.org/10.1021/bi300646y.

88. Ni, Qiang, Ambhighainath Ganesan, Nwe-Nwe Aye-Han, Xinxin Gao, Michael D. Allen, Andre Levchenko, and Jin Zhang. 2011. “Signaling Diversity of PKA Achieved via a Ca2+-cAMP-PKA Oscillatory Circuit.” Nature Chemical Biology 7 (1): 34–40. https://doi.org/10.1038/nchembio.478.

89. Smith, F. Donelson, Jessica L. Esseltine, Patrick J. Nygren, David Veesler, Dominc P. Byrne, Matthias Vonderach, Ilya Strashnov, Claire E. Eyers, Patric A. Eyers, Lorene K. Langberg, John D. Scott. 2017. “Local protein kinase A action proceeds through intact holoenzymes.” Science 356(6344): 1288–1293.

90. Yang, Pei-Chi, Britton W. Boras, Mao-Tsuen Jeng, Steffen S. Docken, Timothy J. Lewis, Andrew D. McCulloch, Robert D. Harvey, and Colleen E. Clancy. 2016. “A Computational Modeling and Simulation Approach to Investigate Mechanisms of Subcellular cAMP Compartmentation.” PLOS Computational Biology 12 (7): e1005005. https://doi.org/10.1371/journal.pcbi.1005005.

91. Zhang, Jin, Christopher J. Hupfeld, Susan S. Taylor, Jerrold M. Olefsky, and Roger Y. Tsien. 2005. “Insulin Disrupts *β*-Adrenergic Signalling to Protein Kinase A in Adipocytes.” Nature 437 (7058): 569–73. https://doi.org/10.1038/nature04140.

92. Zhang, Mingxu, Tommaso Patriarchi, Ivar S. Stein, Hai Qian, Lucas Matt, Minh Nguyen, Yang K. Xiang, and Johannes W. Hell. 2013. “Adenylyl Cyclase Anchoring by a Kinase Anchor Protein AKAP5 (AKAP79/150) Is Important for Postsynaptic *β*-Adrenergic Signaling.” Journal of Biological Chemistry 288 (24): 17918–31. https://doi.org/10.1074/jbc.M112.449462.

